# Hampered AMPK-ULK1 cascade in Alzheimer’s disease (AD) instigates mitochondria dysfunctions and AD-related alterations that are alleviated by metformin

**DOI:** 10.1101/2024.12.28.630598

**Authors:** Arnaud Mary, Samantha Barale, Fanny Eysert, Audrey Valverde, Sandra Lacas-Gervais, Charlotte Bauer, Sabiha Eddarkaoui, Luc Buée, Valérie Buée-Scherrer, Frédéric Checler, Mounia Chami

## Abstract

**Background:** Mitochondrial structure and function alterations are key pathological features in Alzheimer’s disease (AD) brains. The adenosine monophosphate-activated protein kinase (AMPK) and its downstream effector Unc-51 like autophagy activating kinase 1 (ULK1) represent a key node controlling mitochondria health, the alteration of which likely contribute to AD development.

**Methods:** We designed this study to investigate AMPK-ULK1 activation state in post-mortem human sporadic AD brains, in 3xTgAD mice that recapitulate most of human AD features, and in neuronal cells expressing the amyloid precursor protein with the familial Swedish mutation (APPswe). We examined the impact of the pharmacological and genetic modulation of AMPK-ULK1 cascade on mitochondria structure and functions in APPswe cells. We evaluated the potential beneficial impact of AMPK-ULK1 activation by Metformin (Met) on mitochondria defects, as well as on early- and late-stage AD-related alterations *in vivo* and *ex vivo*.

**Results:** At first, we show that AMPK-ULK1 cascade is defective in murine and human AD brains as well as in APPswe cells. We then report that Met administration to 3xTgAD mice alleviates the alterations of neuronal mitochondria structure and function and we consolidate these results in cells using both pharmacological and genetic tools to modulate AMPK-ULK1 cascade. In mice brains, Met reduces the early accumulation of APP C-terminal fragments (APP-CTFs) as well as the amyloid beta (Aβ) burden present in aged mice. Mechanistically, we show that Met increases the localization of APP-CTFs within cathepsin D-positive lysosomal compartments *in vivo* and enhances cathepsin D activity *in vitro*. The reduction of Aβ load by Met occurs through an increased recruitment of Iba1^+^ cells to Aβ plaques and an enhancement of the phagocytic activity of microglia. Accordingly, in symptomatic 3xTgAD mice, Met alleviates microgliosis and astrogliosis, modulates microglia morphology, reduces peripheral proinflammatory cytokines levels, and regulates the expression of a set of inflammatory genes. In addition, Met normalizes dendritic spines shape in organotypic hippocampal slice cultures modeling AD and improves learning performance of 3xTgAD mice.

**Conclusions:** Our study demonstrates potential therapeutic benefits of targeting AMPK-ULK1 cascade to reverse both early and late AD-related alterations, deserving further investigation in fundamental research and in human clinical studies.

## Background

Alzheimer’s disease (AD) is primarily defined by neuronal loss culminating in learning and memory deficits in patients (1). Moreover, AD brains show upstream histopathological alterations tightly linked to disease onset and progression. This comprises microscopic intracellular lesions named neurofibrillary tangles (NFT) formed by the aggregation of hyperphosphorylated forms of the Tau protein, as well as the accumulation of the proteolytic products of the amyloid precursor protein (APP) including the toxic amyloid β (Aβ) peptides forming extracellular senile plaques and the APP-derived C-terminal fragments (APP-CTFs) of 99 amino-acids (C99) and 83 amino acids (C83) (2, 3). Studies converge now to demonstrate that beside Aβ, APP-CTFs are key histopathological features contributing to AD development at an early stage. APP-CTFs accumulate in transgenic and humanized knock-in AD mice models as well as in human sporadic AD brains and iPSCs carrying familial mutations on APP or presenilin 1 encoding genes known to amplify the amyloidogenic processing of APP (4-6). Studies by our group and others demonstrate that APP-CTFs accumulation triggers, upstream to, and independently of Aβ, early lysosomal-autophagic pathology, mitochondrial dysfunctions and mitophagy failure. Notably, APP-CTFs accumulate in intracellular compartments including endosomes, lysosomes and mitochondria in a disease and age-dependent manners (6-11). In parallel to proteinopathy, growing evidence define AD as a consequence of bioenergetics breakdown driven by mitochondrial dysfunction that develops alongside with disease development (12, 13).

Mammalian Adenosine monophosphate-activated protein kinase (AMPK), a highly conserved serine/threonine protein kinase, serves as a master sensor to maintain energy homeostasis (14). AMPK activation controls the biogenesis of new mitochondria and the degradation of defective ones through the phosphorylation of downstream effectors including Unc-51 like autophagy activating kinase 1 (ULK1) (14). While alterations of the status of AMPK phosphorylation was studied in AD, that of ULK1 remains not explored.

This study reports the alteration of AMPK-ULK1 signaling cascade in human sporadic Alzheimer’s brains, and in cellular and mice models of familial AD. Of interest, we unravel the impact of AMPK-ULK1 modulation on mitochondrial structure and function *in vitro* and *in vivo* and on dendritic spines alterations *ex vivo*. Restoring AMPK-ULK1 cascade in AD mice brains reduces proteinopathy linked to APP processing, balances chronic neuroinflammation, and alleviates learning deficit.

## METHODS

### Experimental design

We analyzed the expression of total and active phosphorylation states AMPK and ULK1 in human sporadic Alzheimer’s brains and age-matched heathy individuals and in 10-11 months-old wild-type (WT) and 3xTgAD female mice (Fig. 1A). The activation of the AMPK is one of the primary targets by which the antidiabetic drug Metformin (Met) is believed to act (15). We thus designed preclinical studies including wild-type (WT) and 3xTgAD female mice that were randomly divided into non-treated (H_2_O) and Met treated groups. The study includes a group of 3xTgAD mice aged 9-10 months (Fig. 1B) in which we investigated the impact of Met to alleviate late AD-related alterations. The second group of pre-symptomatic 3xTgAD mice aged 3 months was used to study the impact of Met to prevent the alterations occurring at the early disease stage (Fig. 1C). The experimental endpoint was 1 month of Met treatment. The number of mice per group and that required for each experiment were determined beforehand based on previous extensive experience with the 3xTgAD model and further supported by G power analyses (Fig. 1B, C). Mice were allocated to each experiment in a blinded fashion. When adapted, we used the brain extracts or slices to evaluate distinct AD-related paradigms. Immunofluorescence analyses were carried-out on 3 serial sections. Electron microscopy as well as confocal images were taken blindly. NanoString gene expression analysis were blinded. Outliers were identified using GraphPad Prism version 10 for Windows (GraphPad Software, La Jolla, CA, USA; https://www.graphpad.com). Mice that did not swim or reach the platform the first day of the experiment were excluded from the study design in Fig. 1B. We used murine primary microglia to evaluate the impact of Met on phagocytosis (Fig. 1D). We used murine organotypic hippocampal slice cultures to investigate the effect of Met and CC on the dendritic spine morphology (Fig. 1E). Supplementary experiments were performed in control neuroblastoma cells or expressing APP with the Swedish familial mutation (APPswe). In addition to Met, we used Compound C (CC) (an ATP-competitive AMPK inhibitor) and employ genetic tools to modulate AMPK expression and activity.

**Fig. 1.**
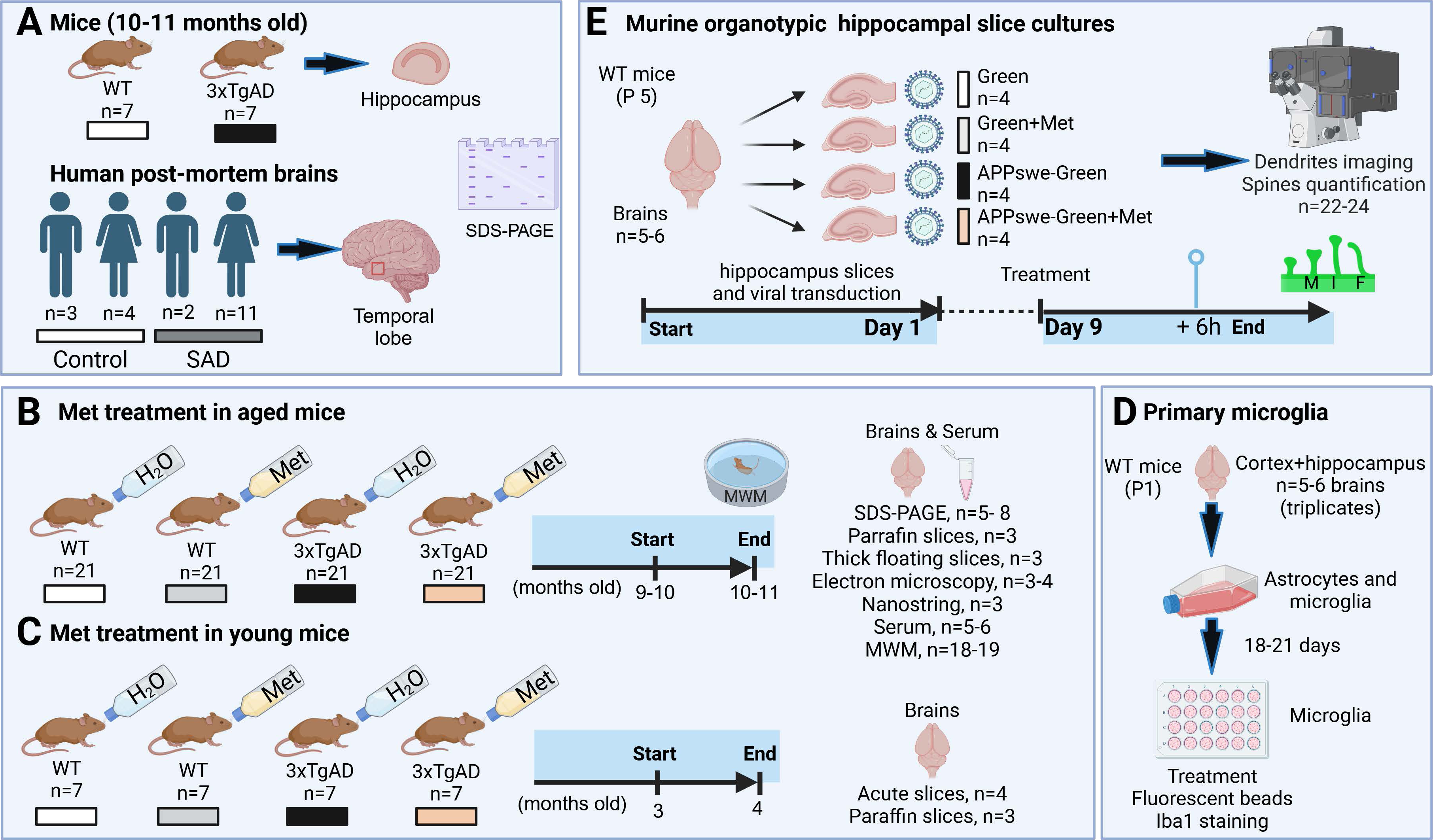
Study Design. **(A)** We analyzed the expression of total and active phosphorylation states AMPK and ULK1 in human sporadic Alzheimer’s brains and age-matched heathy individuals and in 10-11 months-old wild-type (WT) and 3xTgAD female mice when they start to develop Aβ plaques, Tau pathology, and neuroinflammation. (**B**) We studied the impact of Met treatment in symptomatic 3xTg-AD female mice aged 9-10 months. Wild-type (WT) and 3xTgAD female mice were treated for 1 month with Met (2 mg/mL = 400 mg/kg/day) in their drinking water. We performed the Morris Water Maze (MWM) to study the spatial learning and memory. We isolated their brains and used blood sampling for serum collection. This group of mice serves to study mitochondria structure, Aβ and Tau pathologies, and the state of the central as well as peripheral inflammation. (**C**) We investigated the impact of Met treatment (as in (A)) in pre-symptomatic 3xTgAD mice aged 3 months accumulating APP-CTFs with no detectable Aβ, Tau or neuroinflammation. This group of mice serves to study the impact of Met on the early accumulation of APP-CTFs and mitochondrial function in fixed or acute brain slices respectively. (**D**) We used primary microglia to evaluate the impact of Met application (2 mM) for 6 h on phagocytosis. (**E**) We used murine hippocampal slice cultures expressing Green or APPswe-Green that treated or not with Met (2 mM) for 6 h to investigate the effect of Met on the dendritic spine morphology.

### Human brain samples

Studies involving human brains were performed in accordance with the ethical standards of the institutional and/or national research committee and with the 1964 Helsinki declaration and its later amendments or comparable ethical studies. Informed consent for tissue donation for research is obtained by the Brain Bank Neuro-CEB and has been declared at the Ministry of Higher Education and Research (agreement AC-2013-1887) under their approval procedures. We studied control and SAD post-mortem human brains. Cases were anonymized, but information was provided regarding sex, age at death, and neuropathology (Supplementary Table 1).

### Animals

3xTgAD (APPswe: KM670/671NL, TauP301L, and presenilin1 (PS1) M146V) (16) and wild-type (WT; non-transgenic) female mice were housed with a 12:12 h light/dark cycle and were given free access to food and water. All experimental procedures were in accordance with the European Communities Council Directive of 22 September 2010 (2010/63/EU) and approved by the French Ministry of Higher Education and Research (Project number APAFiS#20495-201904231352370) and by Côte d’Azur University Animal Care and Use Committee.

### Mice pharmacological treatment and brains processing

Three-month-old and nine- to ten-month-old 3xTgAD and WT females were treated with metformin hydrochloride (Met) (Merck-Millipore, Cat#317240-5GM) at 2 mg/mL in their drinking water for 1 month (corresponding to ≈ 400 mg/Kg/day, considering 5ml mean daily water consumption that was almost equivalent between the four groups of mice (Supplementary Fig. 1A)). We first report that Met administration did not influence the body weight (Supplementary Fig. 1B). We used the 3xTgAD females that show pronounced AD-related phenotypes versus 3xTgAD males. To note, these later develop the same AD-related alterations but at later age (16). Previous studies have shown that Met rapidly crosses the blood-brain barrier and accumulates in structures of the central nervous system (17). Brains were isolated and snap frozen in liquid nitrogen for SDS-PAGE analyses or incubated in the RNA stabilization reagent (RNAlater, Qiagen, Cat#76106) for 24 h and then stored at -80°C until RNA extraction. Total protein extracts and RNA were obtained from dissected hippocampi. For immunohistochemistry and electron microscopy analyses, mice were anesthetized with a ketamine/xylazine (87 mg/mL and 13 mg/mL, respectively, 1 mL/kg) mixture, transcardially perfused with PBS for 5 min and then with 10 mL of 4% PFA solution (Electron Microscopy Sciences, Cat#15714) for immunohistochemistry or with 10 mL of 2.5% glutaraldehyde (Sigma-Aldrich, Cat#G-5882) in 0.1 M cacodylate buffer for electron microscopy analyses. Brains were further post-fixed overnight at 4°C. For immunohistochemistry, fixed brains were embedded in paraffin and coronal sections were cut with a sliding microtome (8 µm). One mL of blood was collected before the cardiac perfusions and was left to coagulate in a collection tube for two hours at room temperature (RT). Tubes were then centrifuged at 10,000 g, 20 min at 4°C, and the supernatant (serum) was collected, snap-frozen in liquid nitrogen and stored at -80°C until use.

### Morris water maze

Spatial learning and memory tests using the Morris Water Maze (MWM) were performed as previously described (18). Several parameters were evaluated using the Animaze 6.1 Software including the latency to reach the hidden platform (evaluation of learning paradigm), number of target quadrant entry (evaluation of memory paradigm), mean speed (mobility), and latency to visible platform (vision).

### Viral production

We used Green and Green-APPswe constructs cloned in the lentiviral vector with an IRES-ZsGreen fluorescent tag (pHAGE-CMV-MCS-IRES-sGreen, under the control of the CMV promoter (19). Lentiviral particles were produced as previously described by co-transfecting Green or Green-APPswe constructs with two helper plasmids, delta8.9 (packaging vector) and VSV-G (envelope vector) into Lenti-X 293 T cell line (632,180; Clontech, Mountain View, CA, USA) (19). Viral titers were assessed using p24 ELISA (Cell Biolabs, San Diego, Cat#VPK-107).

### Organotypic hippocampal slices preparation and culture

Hippocampal slices were obtained from C57BL/6JRj mice (Janvier Labs, Le Genest Saint-Isle, France) 5 days after birth. Brains were quickly dissected-out to retrieve hippocampi from both hemispheres, sliced onto 400 μm sections and kept into slicing medium (Earles’ Balanced Salt solution (EBSS 97.5%) and EBSS-HEPES (2.5%)). Slices were transferred on sterile hydrophilic membrane millicell discs (Millipore, FHLC01300) placed in semi porous cell culture inserts (Millipore, 0.4 μm) containing culture medium (Minimum Essential Medium Eagle (MEM) + Glutamax-1 (50%), EBSS (18%), EBSS (13%)/D-glucose (5%), penicillin– streptomycin 5,000 U/mL (1%), Horse serum (25%) and Nystatin 10,000 U/mL (0.06%). Two hours after plating, slices were infected with lentiviruses as already described (20). We applied 2 µl of lentiviruses/slice (2 × 10^10^ particles). Slices were kept at 37°C, with 5% CO_2_ for 9 days before the treatment with 2 mM Met or 10µM CC for 6 h and imaging analysis. Control slices were untreated or treated with DMSO.

### Imaging of dendrites and quantitation of spine morphology in organotypic hippocampal slices

Organotypic hippocampal slices attached on the millicell membranes were fixed with PFA 4% for one hour and cover-slipped. Spine morphology was assessed with LSM 780 microscope, Plan-Apochromat 63 × /1.40 Oil DIC M27 lens, zoom 3.0 and pinhole at 52. The quantification of spine structures (filopodia “F”, immature “I”, and mature “M”) was performed using ZEN software.

### DAB and Immunofluorescence staining

Mice brain sections were deparaffined in xylene bath and rehydrated in successive 5 min baths of EtOH 100% (2 times), 90%, and then 70%. Antigens were unmasked in a 90% formic acid (Sigma-Aldrich, Cat#33015-1L-M) bath for 5 min for 82E1 antibody (1/800; IBL America) (recognizing Aβ and C99 but not holo-APP) or for 30 min in citric acid solution pH=6 at 90-100°C (Pressure cooker, Vector Laboratories) for 82E1 and anti-Iba1 (1/1,000, FUJIFILM Wako Chemicals) co-staining, Iba1, GFAP (1/1,000, Novus Biologicals), or AT8 (recognizing pSer202/Thr205-Tau (1/1,000, Thermo Fisher Scientific)) staining. Non-specific binding sites were blocked for 1 h in 5% BSA, 0.05% Triton in PBS solution. Sections were incubated at 4°C overnight with primary antibodies. After three washes with PBS supplemented with 0.02% tween, sections were incubated for one hour with adequate secondary horseradish peroxidase antibodies (Interchim, Montluçon, France) and then revealed with the DAB-ImmPACT system (Vector Laboratories, Cat#PK-6102). Slices were rinsed, counterstained with cresyl violet, and mounted in Isomount 2000 medium (VWR Chemicals). Co-immuno-fluorescent staining was revealed after the incubation for one hour with adequate secondary antibodies coupled with Alexa-488 or Alexa-594 fluorophores (1/1,000, Molecular Probe). Nuclei were stained with DAPI (1/5,000, Invitrogen, Cat#D1306) and slices were mounted with Vectamount medium (Vector, Cat#H-5501-60). Images were taken with a confocal Leica TCS SP5 microscope (Immunofluorescence) or with DMD108 Leica microsystem (DAB) and processed using ImageJ software.

### Microglia morphological analysis

Images of Iba1-stained slices were taken with a confocal Leica TCS SP5 microscope (63X objective, 1 µm-thick Z sections) and processed using ImageJ software as previously described (21). Individual Iba1 positive cells were cropped and thresholded to generate a binary (black and white) image, and the background was manually cleaned. The cell-body area was measured using the freehand selection tool in ImageJ. A Sholl analysis was performed using the ImageJ plugin Sholl Analysis (radius: 4 µm-70 µm, radius step size: 2 µm, radius span: 0 µm). Skeleton analysis was performed on the same binary image, by using the skeletonized using the ImageJ Plugin Skeletonize3D and the plugin AnalyzeSkeleton. The 3D rendering of microglia was obtained using the Imaris software (64X version 9.6.1).

### Transmission electron microscopy

Fixed mice brains were sliced (200 µm) on a vibratome. Two mm^3^ cubes from the subiculum area were micro dissected under binoculars and post-fixed in osmium tetroxide (1% in cacodylate buffer 0.1 M). Brain samples were dehydrated with several incubations in increasing concentrations of ethanol or acetone, respectively, and embedded in epoxy resin (EPON), and 70 nm ultrathin sections were contrasted with uranyl acetate and lead citrate and observed with a Transmission Electron Microscope (JEOL JEM 1400) operating at 100 kV and equipped with an Olympus SIS MORADA camera. We used ImageJ software to analyze mitochondria ultrastructure and to quantify lysosomes.

### Measurements of mitochondrial membrane potential in acute brain slices

Mice were anesthetized with gaseous isoflurane, beheaded, and their brains were collected to be cut into 150 µm thick hippocampal sections on a vibratome. Sections were maintained during 5 h into a solution of artificial cerebrospinal fluid (aCSF: 124 mM NaCl, 3.75 mM KCl, 2 mM CaCl_2_, 1.25 mM NaH_2_PO_4_, 2 mM MgCl_2_, 10 mM D-glucose, 26.5 mM NaHCO_3_, pH=7.43), continuously carbogeneted, at 37°C. Brain sections were then incubated in carbogeneted aCSF with 20 nM tetramethyl rhodamine methyl ester (TMRM, Fluka Analytical, Cat#87919) probe for 30 min at 37°C. TMRM is a cell-permanent dye that accumulates in active mitochondria with intact membrane potentials. Slices were mounted and imaged with a confocal Leica TCS SP5 microscope (63X objective, 1 µm-thick Z sections). The integrated mean fluorescent intensity, reflecting mitochondrial membrane potential was measured in the neuronal soma using ImageJ software.

### Protein extraction and SDS-PAGE analysis

Total protein extracts were obtained from cells or dissected mice hippocampi, using lysis buffer (50 mM Tris pH=8, 10% glycerol, 200 mM NaCl, 0.5% Nonidet p-40, and 0.1 mM EDTA) supplemented with protease inhibitors (Complete, Roche diagnostics). Proteins were resolved in 10% Tris-Glycine SDS-PAGE to reveal total AMPK, p(Thr172)AMPK, total ULK1 and p(Ser555)ULK1. Full-length APP, APP-CTFs, and Aβ were resolved on 16.5% Tris-Tricine SDS-PAGE. Membranes were boiled in PBS, saturated in TBS, 5% skimmed milk, and incubated overnight with specific antibodies. Information about the primary antibodies can be found in Supplementary Table 2.

### Aβ_1-40_ and Aβ_1-42_ and Tau species sandwich ELISAs

The concentrations of human soluble Aβ_1-40_ and Aβ_1-42_ were measured in total protein extract of hippocampi, by using the respective ELISA kits (Invitrogen, Cat#KHB3481 and Cat#KHB3441) as already described (22). The concentrations of phosphorylated human Tau on Thr231, Ser199, and Ser396 residues were similarly determined in total protein extract of hippocampi (diluted at 1:50 in sample diluent buffer) using ELISA kits (Invitrogen, Cat #KHB8051, Cat #KMB7041, Cat #KHB7031), according to the manufacturer’s instructions. The concentrations of human Tau (7C12 N-terminal Tau antibody) and total Tau (TauE7 antibody against prolin rich region) (23). The recombinant Tau (1N4R) was used for the standard curve. Samples were diluted at 1:32 for human Tau and 1:120 for total Tau. Following incubation with an anti-Exon1 antibody or an anti-Tau C-terminal detection antibody, revelation was done with Tetramethyl benzidine (Sigma-Aldrich) and sulfuric acid addition using a spectrophotometer (Multiskan FC, Thermo) at 450 nm.

### Plex Proinflammatory Panel 1 Mouse kit

Levels of interferon (IFN)-γ, interleukin (IL)-1β, IL-2, IL-4, IL-5, IL-6, IL-10, IL-12p70, KC/GRO, and tumor necrosis factor (TNF)-α, were analyzed using the V-PLEX Proinflammatory Panel 1 Mouse Kit (MSD Cat#K15048D), according to the protocol supplied by the manufacturer. The level of IL-4 and IL-12p70 was below the background level in the serum of WT and 3xTgAD mice.

### Primary microglia culture and phagocytosis assay

Primary microglia from WT mice were cultured as previously described (24). Isolated cells were cultured until DIV 21 and then passed in 24-well plaques (2.10^5^ cells/well) containing polyethyleneimine-coated coverslips and cultured for two days. Microglia were treated with Met or CC for 6 h. One µL of fluorescent latex beads (1 µm diameter, Sigma L1030) was pre-opsonized in 4 µL fetal bovine serum for 1 h at 37°C and diluted in DMEM medium (0.01% final beads concentration). The media was replaced with the media containing the beads for 1 h, then the microglia were washed five times with cold 1X PBS and fixed with 4% PFA for 15 min. Microglia were then subjected immunostaining with Iba1 antibody. Cells were observed with a Zeiss Axioplan 2 Fluorescence microscope with a 20X objective. Images were processed with ImageJ to quantify the number of Iba1 positive cells containing at least one bead (phagocytic cell) and the number of beads per Iba1 positive cells.

### *In vitro* Cathepsin D activity

Cell pellets from control as well as APPswe cells (Supplementary Methods and (22)) untreated or treated with Met (2 mM, 6 h) were washed with PBS 1X and centrifuged at 600 g for 5 min. The pellet was resuspended in Tris-HCl (10 mM, pH=7.5). To monitor cathepsin D (Cat D) activity, 40 μg of protein extracts were incubated in acetate buffer (25 mM, pH 4.5, 100 μL and 8 mM L-cysteine-HCl) containing Cat D substrate (Enzo BML-P145-001, 50 µM) in the absence or presence of the 20 µm pepstatin A inhibitor (Sigma, Cat#77170). The fluorochrome was excited at 320 nm and the fluorescence recorded at 420 nm in a fluorescence plate reader (Varioskan, ThermoFisher scientific). The fluorescence was recorded every 30 seconds during 25 min. Specific Cat D activity corresponds to the difference of fluorescence in absence and presence of pepstatin A. Cat D activity is reflected by the slope and the area under the curves.

### NanoString nCounter gene expression

RNA concentration and purity were evaluated by nanodrop analysis. We used the predesigned mouse neuroinflammatory panel (NanoString Technologies, Seattle, WA). Briefly, 100 ng of total RNA was hybridized to capture-reporter probe codesets and immobilized on NanoString cartridges. Excess RNA and probes were removed, and digital barcodes were counted. Raw RCC files were imported to ROSALIND streamlines data analysis following the NanoString guidelines transcripts counting, quality control, and normalization using 13 housekeeping genes (*Aars, Asb10, Ccdc127, Cnot10, Csnk2a2, Fam104a, Gusb, Lars, Mto1, Supt7l, Tada2b, Tbp, Xpnpep1*) and six positive control genes, representing a standard curve of expression abundance. We applied settings for the filter begin with a p value < 0.05 to identify differentially expressed genes.

### Statistical analyses

Data were expressed as means ± SEM. Sample size and replicates (mice, cells, slices) for each experiment are indicated in the Fig. and in captions. Data were analyzed with GraphPad Prism version 10 for Windows (GraphPad Software, La Jolla, CA, USA; https://www.graphpad.com). Data were first analyzed for normal distribution. We used the Mann–Whitney test when the two groups of variables have not passed the normality test. Groups of more than two variables that have passed normality test were analyzed by a 2-Way ANOVA test followed by a Tukey’s multiple comparison test. Groups with less than two variables were analyzed by a one-way ANOVA with Tukey’s multiple comparisons post-test. Kruskal–Wallis test and Dunn’s multiple comparisons post-test was used when groups of variables have not passed the normality test. For MWM, latencies were analyzed using InVivoStat software, by mixed model ANOVA with repeated measures (genotype × treatment × training day) and followed by post-hoc comparisons with Holm correction of the *P*-value for multiplicity. Significant differences are: \**P* < 0.05, \*\**P* < 0.01, \*\*\**P* < 0.001, \*\*\*\**P* < 0.0001 and ns: non-significant. Statistical analyses were reported on signal intensities between fields (Fig.3G, 3K, 3L), Aβ plaque size (Fig. 3H), or between individual cells (Fig. 4D, 4E). In these cases, we also reported in Figure captions the biological effects (Cohen’s d) between mice (treatment effect) using a free online effect size calculator (https://www.socscistatistics.com/effectsize/default3.aspx).

**Fig. 2.**
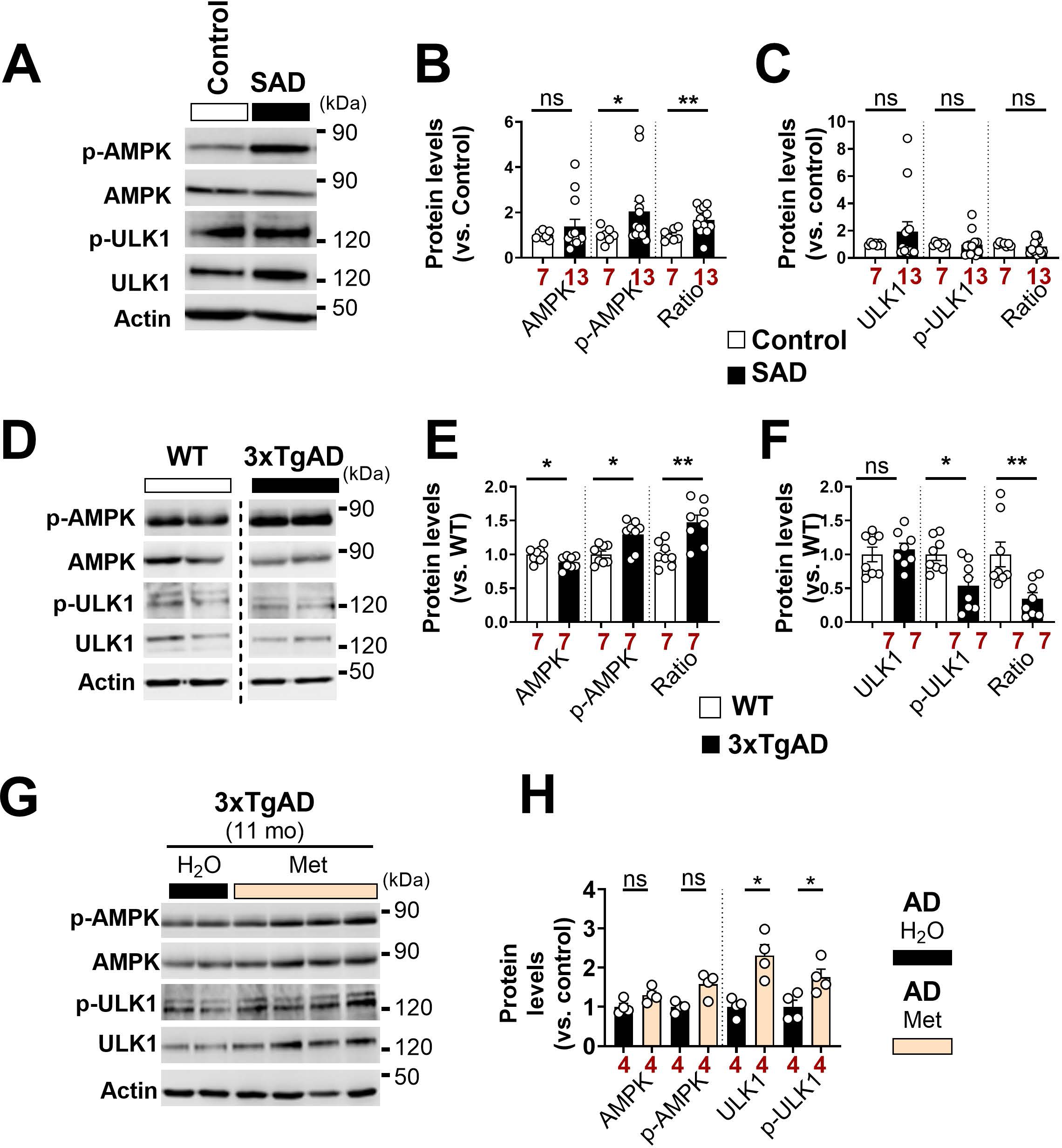
AMPK-ULK1 signaling cascade is altered in post-mortem human AD brains and in hippocampi of symptomatic 3xTgAD mice. (**A, D**) Representative SDS-PAGE, and (**B, C, E, F**) the quantitative graphs showing protein levels of p(Thr172)AMPK, total AMPK, p(Ser555)ULK1, and total ULK1 and the ratio of the phosphorylated forms of AMPK and ULK1 versus respective total proteins in the total homogenates of temporal lobe of AD (n=13) and control (n=7) brains (A-C) (patients information in Supplementary Table 1) and in hippocampi of Wild-type (n=7) and 3xTgAD mice aged 10-11 months (n=7) (D-F). Actin was used as loading control. (**G**) Representative SDS-PAGE and (**H**) the quantitative graphs of p(Thr172)AMPK, total AMPK, p(Ser555)ULK1, and total ULK1, protein levels and the ratio of the phosphorylated forms of AMPK, and ULK1 versus respective total proteins in the hippocampi of 11-month-old 3xTgAD mice untreated (H_2_O) (n = 4) or treated with Met (2mg/ml) in the drinking water (n = 4) for one month. Actin was used as loading control. Graphs represent means ± S.E.M versus Control or WT mice (taken as 1). * *P* <0.05, ** *P* <0.01 and ns: non-significant using Mann Whitney test.

**Fig. 3.**
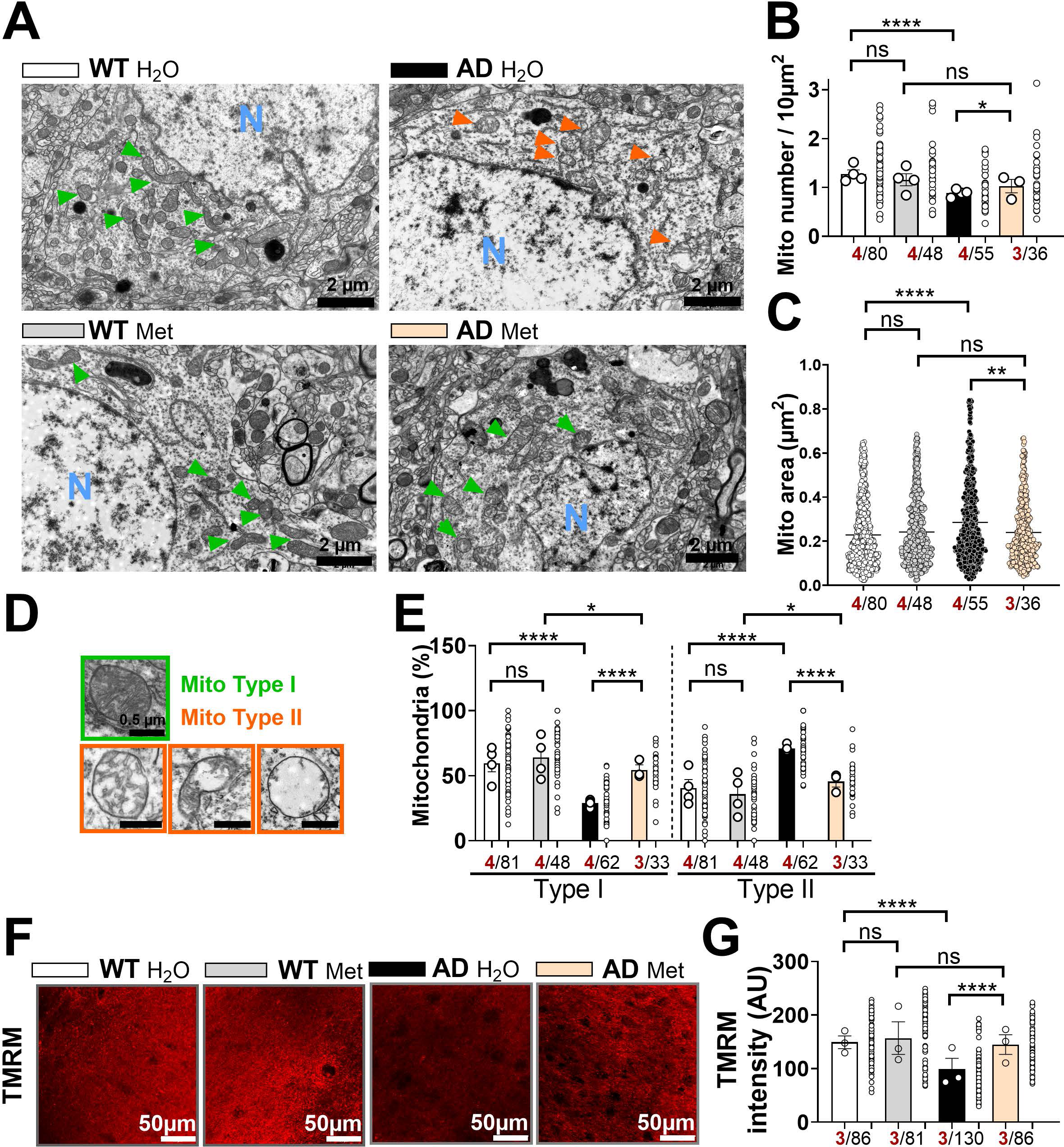
Metformin alleviates mitochondrial structure and function defects *in vivo*. (**A**) Representative electron microphotographs of neuronal soma of wild type (WT) and 3xTgAD mice aged 10-11 months untreated (H_2_O) or treated with (Met). N: nucleus. Scale bars, 2 µm. (**B**, **C**) Quantitative graphs of the mitochondria number/10 µm^2^ (B) and mitochondria area (µm^2^) (C). (**D**) Representative microphotographs of mitochondria type I and type II (Green and orange arrowheads in (A). Scale bars, 0.5 µm. (**E**) Quantitative graphs of mitochondria types distribution. (**F**) Representative TMRM staining obtained in acute brain slices in the subiculum area of WT and 3xTgAD mice aged 4 months treated as in (A). Scale bars, 50 µm. (**G**) Quantitative graph of TMRM intensity in neuronal soma. All data are presented as mean ± SEM. The number of mice in each condition is indicated in red. The number of analyzed fields (B), single mitochondria (C and E), and Soma (G) is indicated in black. *P* values were calculated using a 2-Way ANOVA test followed by a Tukey’s multiple comparison test. * *P* <0.05, ** *P* <0.01, **** *P* <0.0001, and ns: non-significant.

**Fig. 4.**
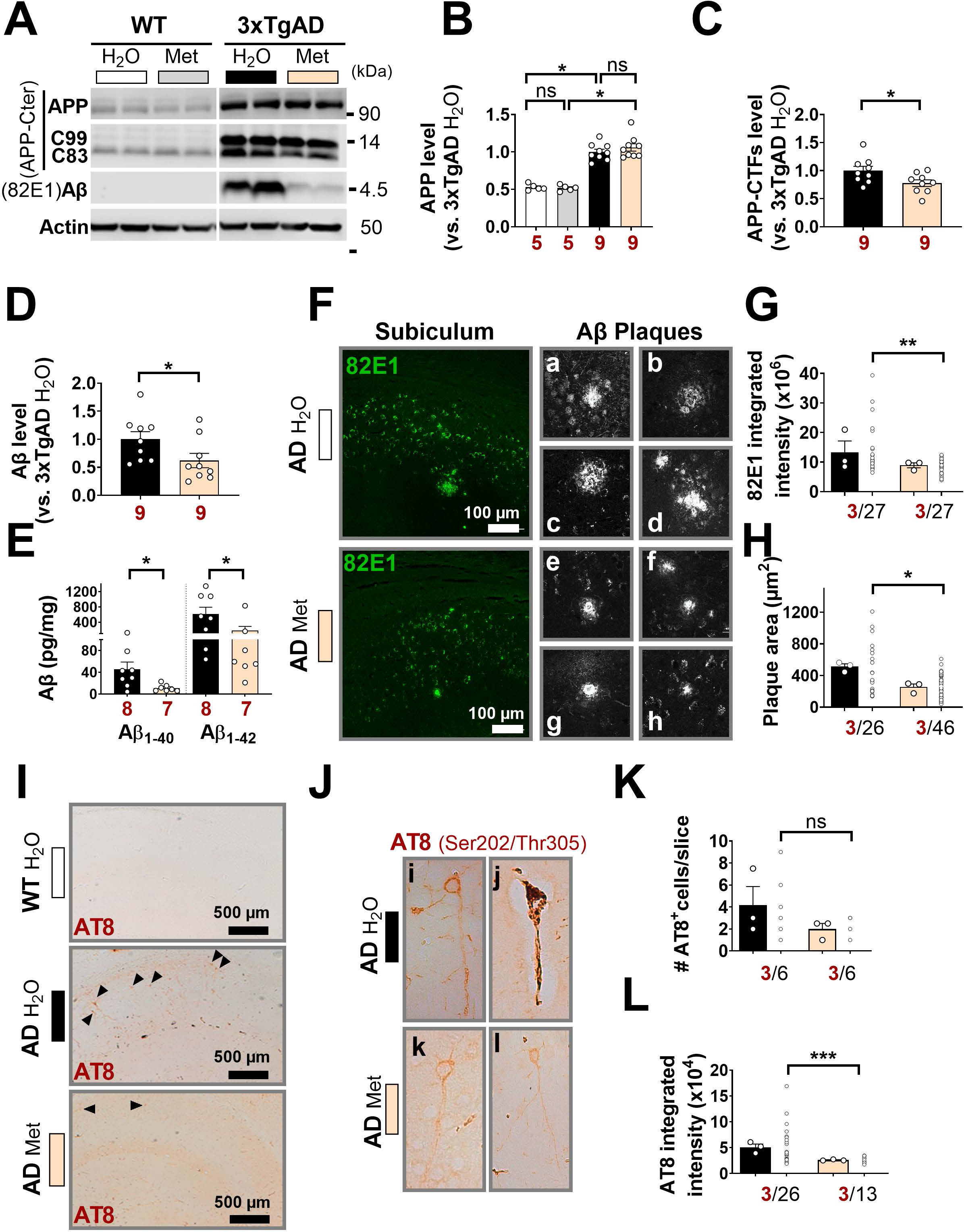
Metformin reduces β amyloid in 3xTgAD mice. (**A**) Representative SDS-PAGE, and (**B**-**D**) Quantitative graphs showing full-length APP (APP) (B), APP-CTFs (i.e. C99 and C83) (C), and Aβ protein levels (D) in the hippocampi of wild type (WT) and 3xTgAD (AD) mice aged 10-11 months treated as in Fig. 1A. Actin was used as loading control. (**E**) Levels of Aβ_1-40_ and Aβ_1-42_ (pg/mg of total proteins) detected by ELISA in the hippocampi total protein homogenates. (**F**) Representative images showing 82E1 fluorescent signal in the hippocampus (subiculum area) and of amyloid plaques in AD H_2_O (a, b, c, d) and in AD Met (e, f, g, h). Scale bars, 100 µm. (**G**, **H**) Quantitative graphs of 82E1 integrated intensity (G) and of Aβ plaque area (µm^2^). (**I**) Representative images showing AT8 signal (recognizing Ser202/Thr205 Tau epitopes) in the hippocampi (subiculum area). Scale bars, 500 µm. (**J**) Representative images showing AT8 signal in individual neurons of AD H_2_O (i, j) and of AD Met (k, l). (**K**, **L**) Quantitative graphs of AT8^+^ cells (K) and of AT8 integrated signal intensity (L). All data are presented as mean ± SEM. The number of mice is indicated in red. The number of analyzed slices (G, K), Aβ plaques (H) and AT8^+^ cells (L) is indicated in black. *P* values were calculated using Mann Whitney test. * *P* <0.05, ** *P* <0.01, *** *P* <0.001, and ns: non-significant. The effect size between mice Cohen’s *d* (0.894338 (G), 3.984033 (H), 1.003044 (K) and 3.220608 (L)).

## RESULTS

### AMPK-ULK1 cascade is hampered in human Alzheimer’s brains and in murine and cellular AD models

We examined the expression of total and phosphorylated AMPK and ULK1 proteins in human sporadic Alzheimer’s brains and in age-matched controls (Supplementary Table 1 and Fig. 1A) and in the hippocampi of 3xTgAD mice and their control littermates at 10-11 months-old (Fig. 1D). We report an enhancement of p(Thr172)AMPK expression level and of p(Thr172)AMPK/total AMPK expression ratio in human AD and 3xTgAD brains versus control/WT brains (Fig. 2A, B, D, E). While the level of total AMPK remained unchanged in human AD brains (Fig. 2A, B), it is reduced in 3xTgAD brains (Fig.2 D, E). Intriguingly, despite higher AMPK active phosphorylation, human AD brains show unchanged levels of total ULK1 and of its active phosphorylated form p(Ser555)ULK1 (Fig. 2A, C). We consistently observed a reduction of p(Ser555)ULK1 expression level and of p(Ser555)ULK1/total ULK1 ratio in the 3xTgAD hippocampi (Fig. 2D, F). These results suggest a blockade of AMPK-ULK1 signaling cascade in human and murine AD brains. We also examined the status of AMPK-ULK1 cascade in the neuroblastoma cells expressing APPswe (Supplementary Fig. 2A). We first validated the overexpression of APPswe and reported enhanced APP-CTFs (C83 and C99) accumulation compared to control cells (Supplementary Fig. 2A). Surprisingly, we reported a drastic reduction of p(Thr172)AMPK expression level and of p(Thr172)AMPK/total AMPK ratio in APPswe expressing cells (Supplementary Fig. 2A, B). This was associated with a reduction of p(Ser555)ULK1 expression level and of p(Ser555)ULK1/total ULK1 ratio (Supplementary Fig. 2A, C).

Together, these data converge to show a blockade of AMPK-ULK1 signalling cascade in human sporadic AD brains, and in cellular and mice AD models mimicking familial AD forms.

### Metformin restores the alteration of mitochondria structure and function in 3xTgAD mice brains and APPswe cells

We designed a first set of experiments including the 3xTgAD (AD) mice aged 9-10 months and age-matched control wild type (WT) mice that were untreated (H_2_O) or treated with metformin (Met) (2mg/ml in the drinking water for 1 month) (Fig. 1B). We validated the activation of AMPK-ULK1 cascade by Met through an enhancement of p(Ser555)ULK1 in the hippocampi of Met-treated 3xTgAD mice (Fig. 2G, H). Intriguingly, we also observed an increase of the expression of total ULK1 in 3xTgAD mice treated with Met (Fig. 2G, H). We studied the impact of Met on mitochondria ultrastructure by electron microscopy (EM) and analysed mitochondria number, morphology and size in the subiculum area of untreated and treated WT and 3xTgAD mice. As we previously reported (6), we observed a reduction of mitochondrial number (Fig. 3A, B), as well as an increase of mitochondria area (Fig. 3A, C) in the neuronal somas of 3xTgAD untreated mice vs WT untreated mice (Fig. 3A-C). Met treatment reverses these alterations in 3xTgAD neuronal soma by enhancing mitochondria number and reducing their area enlargement, while having no effect in WT mice (Fig. 3A-C). We then identified two mitochondria morphological types, were type I corresponds to mitochondria with uniform matrix filled with densely packed and regularly distributed cristae, and type II corresponds to mitochondria showing morphological abnormalities, ranging from focal loss of cristae with empty spaces to severe loss of cristae and reduced matrix density (Fig. 3D). The quantification revealed a reduction of mitochondria type I and an increase of mitochondria type II in untreated 3xTgAD mice neurons (Fig. 3A, D, E). The treatment with Met significantly shifted mitochondria shape towards type I in 3xTgAD treated mice while having no effect in WT mice (Fig. 3A, D, E).

We quantified mitochondrial membrane potential as a read-out of mitochondrial functional state *in vivo*. These analyses were performed in acute brain slices prepared from WT and 3xTgAD mice untreated or treated with Met (Fig. 1C). We used TMRM probe, the loss in intensity of which reflects mitochondrial membrane depolarization. We report a reduction of TMRM signal in the neuronal somas of 3xTgAD mice hippocampi as compared to WT mice (Fig. 3F, G). Met treatment preserved mitochondrial membrane potential in 3xTgAD mice hippocampi and had no effect in WT mice (Fig. 3F, G).

We then sought to study the impact of the pharmacological and genetic targeting (blockade and activation) of AMPK in APPswe-expressing cells. We first show, as expected, that Met enhanced p(Thr172)AMPK and p(Ser555)ULK1 levels and their ratios versus total AMPK and total ULK1, respectively(Supplementary Figure 2D-F). We also report that pharmacological AMPK inhibition with compound C (CC) triggers, as expected, a repression of AMPK-ULK1 cascade (reduced p(Thr172)AMPK and p(Ser555)ULK1 and their ratios versus total AMPK and total ULK1, respectively) (Supplementary Figure 2G-I). We previously reported that APPswe-expressing cells exhibit reduced number of mitochondria, characterized by enhanced size respective to their controls (6). We investigated the impact of Met and CC treatments on mitochondrial ultrastructure and function in APPswe cells treated with Met and CC. EM analyses revealed that CC enhances mitochondria number in APPswe cells as compared to vehicle-treated cells (Supplementary Fig. 3A, B) associated with a drastic fragmentation of mitochondria (reduction of mitochondria area) (Supplementary Fig. 3A, C). Met treatment significantly enhanced mitochondria number (Supplementary Fig. 3D, E) and significantly but moderately reduced mitochondria area (Supplementary Fig. 3D, F). We previously reported that mitochondrial membrane potential (Δψmit) is reduced and mitochondrial ROS (mitROS) levels are enhanced in APPswe expressing cells (6). We show herein that while CC treatment triggers Δψmit hyperpolarization and exacerbates mitROS levels (Supplementary Fig. 3G, H), Met restored Δψmit to a level similar to that in control cells and did not rescue mitROS (Supplementary Fig. 3G, H). We also report enhanced mitROS and Δψmit in control cells treated with CC but not in those treated with Met (Supplementary Figure 4A, B), thus confirming the deleterious effect of AMPK blockade on mitochondria function in physiological and AD conditions. We validated these findings using lentiviral tools to express a constitutive-active (CA-AMPK: AMPKα 1-312), or a dominant-negative (DN-AMPK: AMPKα K45R) AMPK constructs (Supplementary Figure 5A-C). First and as expected, we show that the expression of CA-AMPK enhances p(Thr172)AMPK expression level and of p(Thr172)AMPK/total AMPK ratio versus APPswe cells expressing AMPKα wild type subunit (WT-AMPK) (Supplementary Fig. 5A, B). On the contrary, the expression of the DN-AMPK triggers a significant reduction of p(Ser555)ULK1 expression level and of p(Ser555)ULK1/total ULK1 ratio (Supplementary Fig. 5A, C). Importantly, the expression of DN-AMPK enhances both Δψmit and mitROS levels, as does CC, but to a lesser extent (Supplementary Fig. 5D, E). CA-AMPK expression partially reduce mitROS levels without affecting Δψmit (Supplementary Fig. 5 D, E).

Together, these data demonstrate that while the activation of AMPK-ULK1 cascade alleviates mitochondrial defects, its blockade exacerbates mitochondrial alterations.

### Metformin ameliorates Aβ pathology in 3xTgAD mice

Since we unraveled beneficial effects associated with AMPK activation in AD cells and most importantly in the 3xTgAD mice, we thought to investigate the impact of AMPK activation on AD-related alterations *in vivo*.

We used the same four groups of mice aged 10-11 months (Fig. 1B) to study the impact of Met treatment on APP processing (Fig. 4A-H) and quantified the number of neurofibrillary tangles (Fig. 4I-L) as well as Tau phosphorylation status (Supplementary Fig. 6). SDS-PAGE analyses showed that Met significantly reduced the levels of APP-CTFs (Fig. 4A, C) and of total Aβ peptides (Fig. 4A, D) in the subiculum of 3xTgAD mice without affecting the expression level of the full-length APP (Fig. 4A, B). Accordingly, ELISA assays reveal a reduction of the level of both Aβ_1-40_ and Aβ_1-42_ peptides in the hippocampi of 3xTgAD mice treated with Met (Fig. 4E). We quantified APP-derived fragments accumulation by immunofluorescence using 82E1 antibody specifically directed towards the N-terminal sequence of soluble and insoluble Aβ peptide and C99 fragment, with no cross-reaction with holo-APP (Fig. 4F-H). 3xTgAD mice treated with Met show reduced 82E1 integrated signal intensity in the subiculum area of the hippocampus (Fig. 4F, G). In addition, Met reduces Aβ plaque area (Fig. 4F, H) as it is shown in representative images of Aβ plaques cropped from the subiculum of untreated (Fig. 4F (a-d)) and Met-treated (Fig. 4F (e-h)) 3xTgAD brain slices. We used adjacent slices and applied immunohistochemistry using AT8 antibody recognizing phosphorylated Tau at Ser202 and Thr205 residues. We quantified the number of AT8 positive cells (Fig. 4I-K), and AT8 integrated intensity in the subiculum (Fig. 4I, J, L). As expected, the number of AT8^+^ neurons and the intensity of AT8 signal were higher in 3xTgAD mice hippocampi versus WT mice (Fig. 4I, J). Met treatment showed a non-significant trend towards a reduction of the number of AT8^+^ positive cells (Fig. 4J, K), as well as a significant reduction of AT8 integrated intensity (Fig. 4J, L) in 3xTgAD mice brains. However, we did not report any impact of Met treatment on Tau phosphorylation at Ser199, Thr231 and Ser396 residues as quantified versus total murine and human Tau (tTau) (Supplementary Fig. 6A), or human Tau (hTau) (Supplementary Fig. 6B) in 3xTgAD hippocampi compared to untreated 3xTgAD hippocampi.

These results demonstrate that Met treatment alleviates AD-related proteinopathy in 10-11 month-aged 3xTgAD mice with a significant impact on APP fragments-associated pathologies.

### Metformin prevents early APP-CTFs accumulation in 3xTgAD mice

We previously reported that APP-CTFs accumulates in the hippocampus of young 3xTgAD mice prior to Aβ intraneuronal detection and plaques Aβ formation (11). Notably, we showed that this early accumulation of APP-CTFs contributes, in an Aβ-independent manner, to AD-related alterations (8, 11). We specifically investigate the impact of Met on the early proteopathy linked to the accumulation of APP-CTFs in the absence of Aβ in a second set of experiments using the 3xTgAD mice aged 3 months (Fig. 1C). As in Fig. 4F, we used immunofluorescence and quantified 82E1 signal intensity reflecting only APP-CTFs load in the hippocampi of young 3xTgAD mice (Fig. 5A, B). We report a reduction of 82E1 integrated signal intensity in the subiculum area of the hippocampus of Met-treated 3xTgAD mice versus untreated mice (Fig. 5A, B). APP-CTFs accumulation was previously shown to hamper lysosomal degradation and this, in turn, contributed to APP-CTFs overaccumulation (11). We quantified the extent of the colocalization of 82E1 signal within lysosomal compartment stained with cathepsin D (Cat D) antibody in the group of 3xTgAD mice aged 3 months (Fig. 1C). We first confirm the presence of a dense 82E1 signal reflecting APP-CTFs intra-organellar accumulation in untreated 3xTgAD neurons (Fig. 5C). Consistently, we report a reduction of 82E1 signal and reveal an enhancement of the colocalization of 82E1 and Cat D signals in Met-treated 3xTgAD mice (visible as yellow merged signal in the representative images of neuronal soma cropped from the hippocampus) (Fig. 5C, D). We then quantitatively demonstrate an increase of the number of Cat D puncta in the neuronal soma of Met-treated 3xTgAD hippocampi versus untreated mice (Fig. 5C, E).

**Fig. 5.**
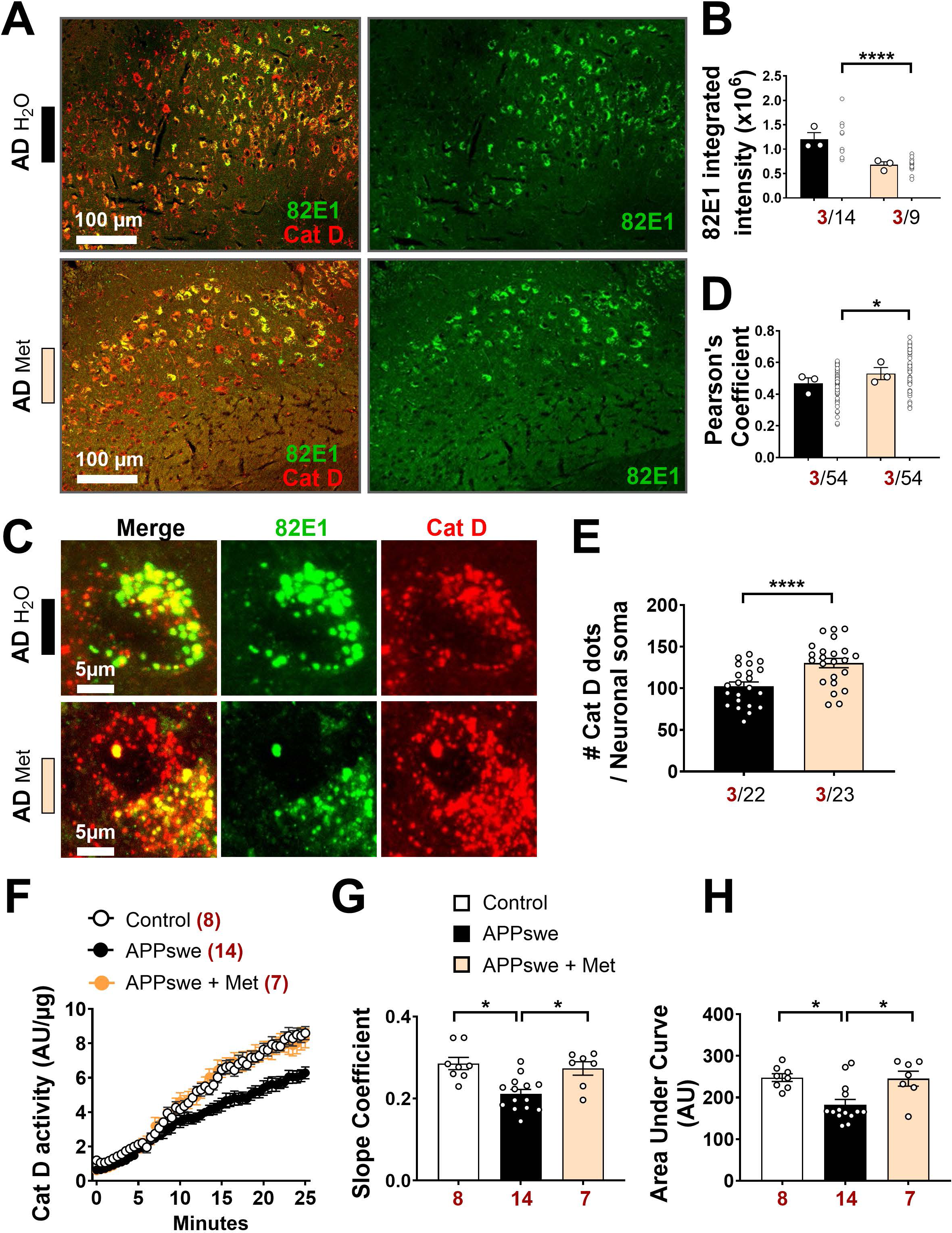
Metformin treatment reduces early APP-CTFs accumulation in 3xTgAD mice. (**A**) Representative images showing 82E1 and Cat D signals in the hippocampi (subiculum area) of 4-month-old 3xTgAD mice untreated (H_2_O) or treated with Met. Scale bars, 100 µm. (**B**) Quantitative graphs of 82E1 integrated signal intensity. (**C**) Representative images showing 82E1 and Cat D signals in the neuronal soma. Scale bars, 5 µm. (**D**, **E**) Quantitative graphs of 82E1-Cat D Pearson’s colocalization coefficient (D), and of the number of Cat D dots per neuronal soma (E). (**F**) Cathepsin D specific activity in Neuroblastoma cells expressing empty vector (Control) or APPswe construct (APPswe) treated or not with Met. (**G, H**) Quantitative graphs of the slope coefficients (G), and Area Under the Curve (H). All data are presented as mean ± SEM. The number of mice (B, D, E) and replicates (G, H) is indicated in red. The number of analyzed slices (B) and neuronal soma (D, E) is indicated in black. *P* values were calculated using Mann Whitney test (B, D, E), or Kruskal-Wallis test and Dunn’s multiple comparisons post-test (G, H). * *P* <0.05, **** *P* <0.0001. The effect size between mice Cohen’s *d* (2.970202 (B) and 1.051959 (D)).

The enhanced number of Cat D^+^ dots/lysosomes may reflect an increase of lysosomal degradation activity. We investigated the impact of Met treatment on Cat D activity and reveal that the specific activity of Cat D is reduced in SH-SY5Y cells expressing APPswe versus controls as demonstrated by reduced slope coefficient (Fig. 5F, G) and area under the curve (Fig. 5F, H). Importantly, Met treatment corrects the loss of Cat D activity in APPswe cells to a level similar to that observed in control cells (Fig. 5F-H).

All together, these data demonstrate that Met treatment reduces early APP-CTFs accumulation in 3xTgAD mice hippocampi likely through enhanced lysosomal degradation.

### Metformin reduces gliosis in aged 3xTgAD mice

In AD, chronic inflammation involves a deregulation of both the central and the peripheral immune systems and is thought to be driven by proteinopathy and bioenergetic dysfunctions (13). We investigated the impact of Met treatment on neuroinflammation mediated by microglia and astrocytes in WT and 3xTgAD mice aged 10-11 months (Fig. 1B). By using immunohistochemistry and Iba1 antibody recognizing microglia and macrophages, we reveal gliosis in untreated 3xTgAD mice aged 10-11 months as indicated by an increase of the total number of Iba1^+^ cells/mm^2^ (Fig. 6A, B). We specifically report an increase of Iba1^+^ “active-like” cells characterized by reduced branching and a dark staining of nuclei (Fig. 6C). We then showed that Met treatment significantly reduces the total number of Iba1^+^ cells and of Iba1^+^ “active-like” and “resting” cells identified with highly ramified morphology in 3xTgAD mice hippocampi, while having no effect in WT mice (Fig. 6A-C).

**Fig. 6.**
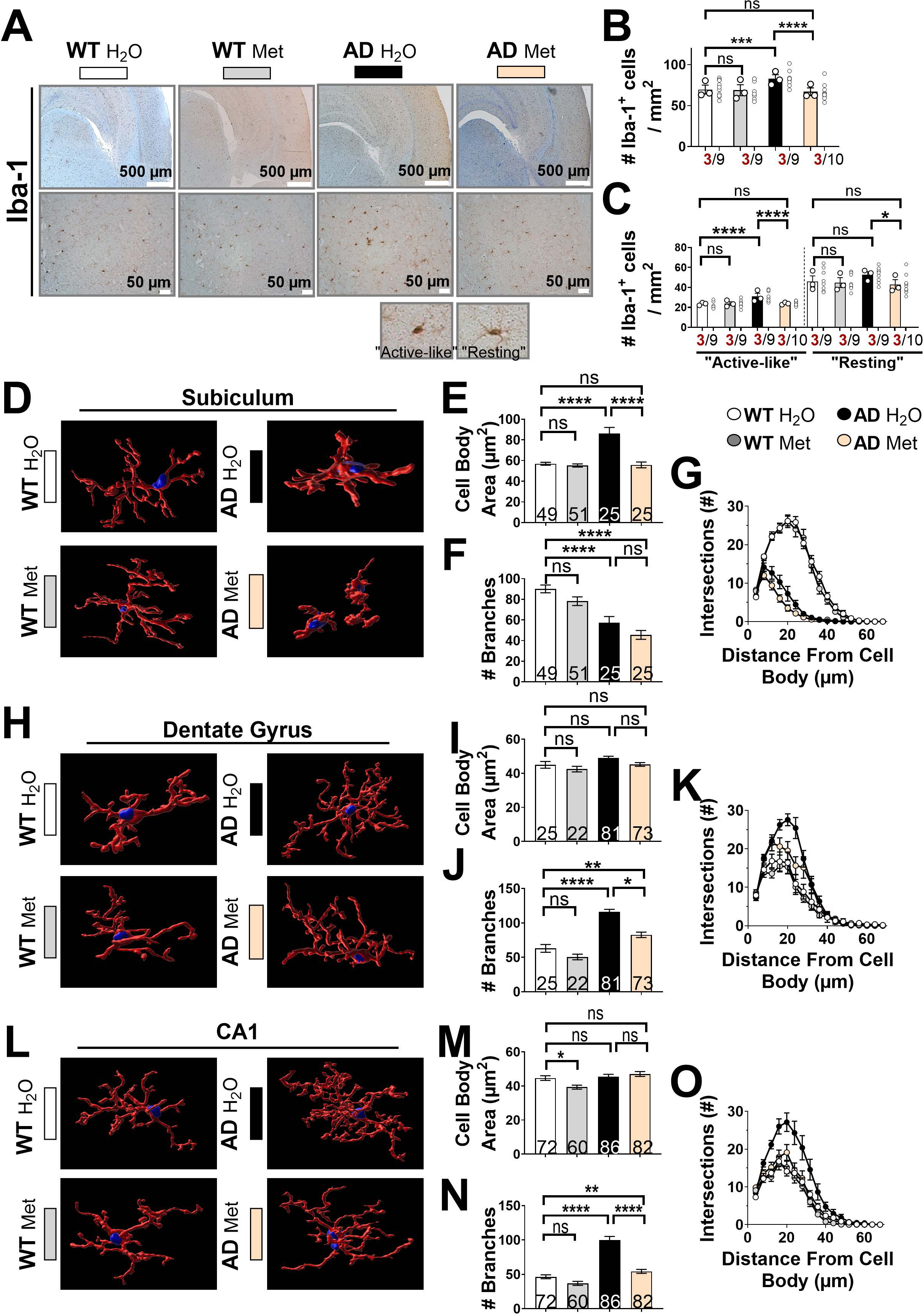
Metformin alleviates gliosis in 3xTgAD mice. (**A**) Representative images showing Iba1 staining in the hippocampus and higher magnification of the subiculum area of 11-month-old WT and 3xTgAD mice untreated (H_2_O) or treated with Met. Scale bars, 500 µm (top) or 50 µm (bottom). (**B**, **C**) Quantitative graphs of Iba1 positive cells (Iba1^+^) (B) and of “active-like” and “resting” Iba1^+^ cells (shown in representative images) (C). (**D**-**O**) Representative 3D-reconstructed Iba1^+^ cells near Aβ plaques in the subiculum area (D), or far from Aβ plaques in the Dentate Gyrus (H) and CA1 (L) regions. Quantitative graphs of the cell body area (µm^2^) (E, I, M), the number of microglia branches (F, J, N), and the number of intersections depending on the distance from the cell body, obtained by the Sholl Analysis (G, K, O). All data are presented as mean ± SEM. The number of mice (B, C) is indicated in red. The number of analyzed slices (B, C) and microglia (E, F, I, J, M, N) is indicated in black. *P* values were calculated using a 2-Way ANOVA test followed by a Tukey’s multiple comparison test. * *P* <0.05, ** *P* <0.01, *** *P* <0.001, **** *P* <0.0001, and ns: non-significant.

To study in depth microglia morphology, we applied high-resolution confocal imaging and analysed 3D-reconstructed Iba1^+^ cells in thick brain slices. We first reveal altered microglia/macrophages morphologies near (subiculum) and far (Dentate Gyrus (DG) or CA1) from Aβ plaques (Fig. 6D-O). We show that Aβ plaque-associated microglia/phagocytes in the subiculum area are characterized by enlarged cell bodies (Fig. 5D-E) and by a drastic reduction of the number of branches (Fig. 6D, F), as well as of the average number of intersections away from cell bodies (Fig.6D, G). However, while microglia/macrophages in the DG and CA1 regions showed unchanged cell body area (Fig. 6 H, I, L, M), they exhibited hyper-ramified skeleton (Fig. 6J, N) and enhanced intersections at each distance from cell bodies (Fig. 6K, O). Interestingly, Met treatment restores microglia/macrophages morphologies near (subiculum) and far (DG and CA1) from Aβ plaques, by reducing cell body area in the subiculum (Fig. 6D, E) and reversing morphological hyper-branching in the DG and CA1 regions (Fig. 6H, J, K, L, N, O).

In addition, we reveal enhanced astrogliosis (increase of the number of GFAP^+^ cells (astrocytes)) in untreated 3xTgAD brains aged 10-11 months compared to age-matched WT mice (Supplementary Fig. 7A, B). We similarly show that Met treatment reduces the number of GFAP^+^ cells in the hippocampi of 3xTgAD mice, while having no effect in WT mice (Supplementary Fig. 7A, B).

### Metformin enhances the recruitment of microglia to Aβ plaques and modulates central and peripheral inflammation

Aβ peptides deposition in senile plaques instigates the recruitment of microglia/macrophages for plaques engulfment and degradation. As the disease advances, microglia morphology changes and shift toward chronic inflammatory responses mediated by the expression of inflammatory markers such as CD68 (25). We applied high-resolution confocal imaging and 3-D reconstruction and quantified Iba1 signal within amyloid plaques (Fig. 7A). We used the group of 3xTgAD mice aged 10-11 months treated or not with Met (Fig. 1B). We reveal an increase of the recruitment of Iba1^+^ cells to Aβ plaques in 3xTgAD mice treated with Met as compared to untreated mice (Fig. 7A, B). We also report a drop of plaque-associated CD68 signal in the 3xTgAD mice treated with Met likely suggesting reduced inflammatory state of recruited microglia and enhanced phagocytic activity (Fig. 7C, D). Accordingly, we demonstrate that Met enhances phagocytic activity of murine primary cultured microglia incubated with fluorescent microbeads as revealed by an increase of the number of microglia with at least one fluorescent bead (Fig. 7E, F) and of the number of beads per microglia (Fig. 7E, G). On the contrary, CC treatment reduces phagocytic activity of murine primary cultured microglia incubated with fluorescent microbeads (Supplementary Fig. 8A-C). These results strongly suggest an anti-inflammatory effect of actived AMPK-ULK1 cascade and support our observed reduced Aβ plaques size in 3xTgAD mice treated with Met (Fig. 4F, H).

**Fig. 7.**
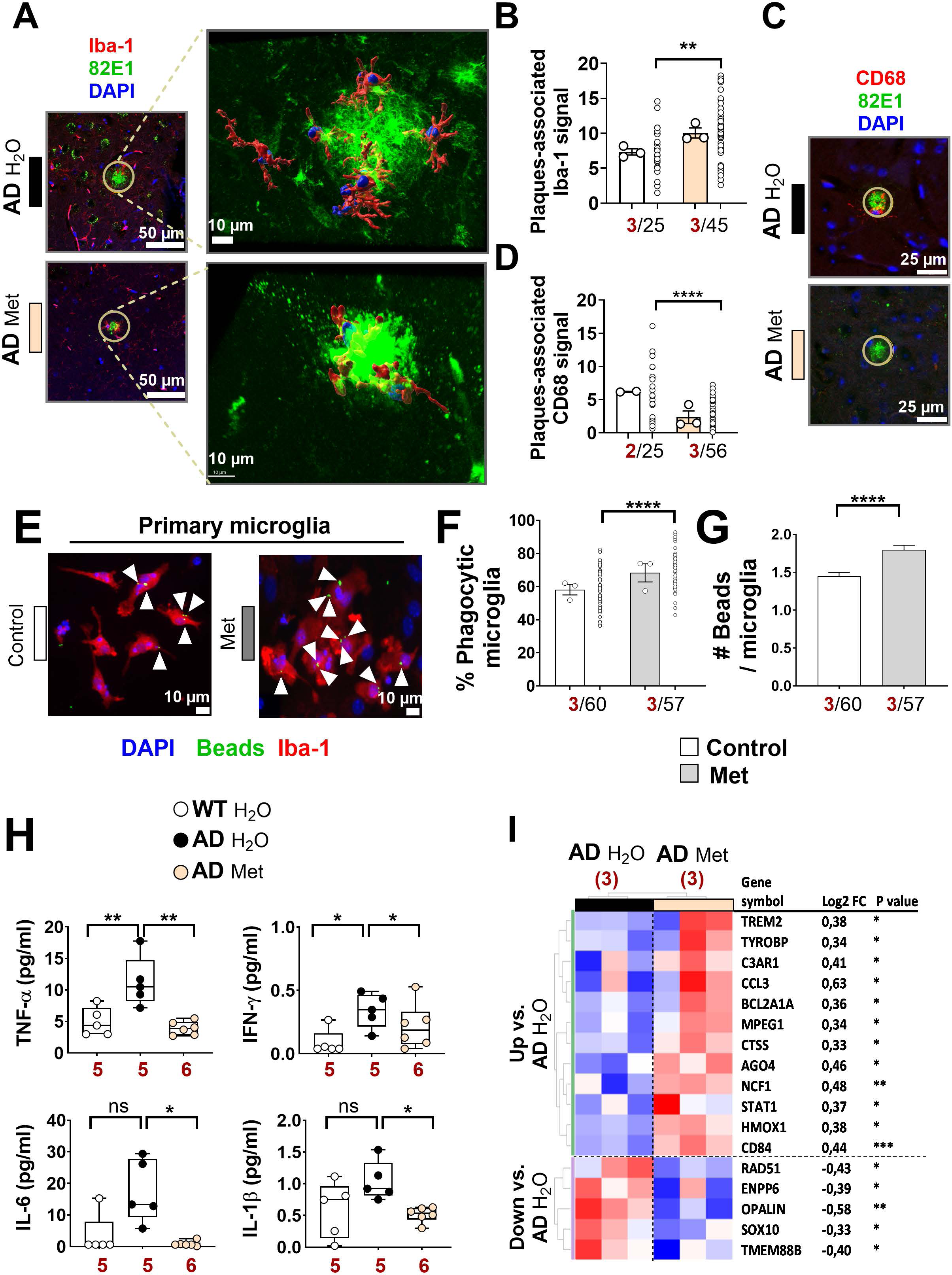
Metformin enhances the recruitment of microglia to Aβ plaques and modulates central and peripheral inflammation. (**A**) Representative images showing 82E1 and Iba1 co-immunostaining in the hippocampus of 3xTgAD mice untreated (H_2_O) or treated with Met. The yellow circle (25µm radius) indicates plaque associated Iba1^+^ cells (left, scale bars 50 µm). The 3D reconstruction of the magnification of Aβ plaques and the Iba1^+^ cells is shown in the right (scale bars 10 µm). (**B**) Quantitative graph of Iba1 signal area in <25 µm radius area. (**C**) Representative images showing 82E1 and CD68 co-immunostaining in the hippocampi of 3xTgAD mice untreated (H_2_O) or treated with Met. (**D**) Quantitative graph of CD68 signal area in <25 µm radius area. (**E**) Representative images of primary microglia incubated with fluorescent microbeads (Green), treated or not with Met and stained with Iba1 antibody (Red). Nuclei were stained with DAPI. Scale bars, 10 µm. (**F**) Quantitative graph of the proportion of phagocytic microglia (that contains at least one microbead). (**G**) Quantitative graph of the number of microbeads per microglia. (**H**) Plots representing the levels of inflammatory markers (TNF-α, IFN-γ, IL-6, IL1-β) (pg/ml) in the serum of WT and 3xTgAD mice untreated (H_2_O) or treated with Met. (**I**) Heat map representing differentially expressed genes (obtained by NanoString) in the hippocampi of 3xTgAD mice untreated (H_2_O) or treated with Met with a cut off of *P* <0.05. Gene symbol and Log2FC (fold change) are indicated in the right. All data are presented as mean ± SEM. The number of mice (B, D, H, I) and the number of replicates (F, G) is indicated in red. The number of analyzed plaques (B, D) and microglia (F, G) is indicated in black. *P* values were calculated using a Mann Whitney test (B, D, F, G), or 1-Way ANOVA followed by a Tukey’s multiple comparisons post-test (H). * *P* <0.05, ** *P* <0.01, **** *P* <0.0001 and ns: non-significant. The effect size between mice Cohen’s *d* (2.597605 (B) and not-determined (D)).

We then examined cytokine protein profiles in the serum of WT and 3xTgAD mice aged 10-11 months treated or not with Met (Fig. 1B) as a read-out of peripheral inflammation. We first report a significant enhancement of proinflammatory cytokines TNF-α, INF-γ levels, and an increase tendency for IL-6 and IL-1β levels in the serum of untreated 3xTgAD versus age-matched WT mice (Fig. 7H). Importantly, Met treatment significantly reduces TNF-α, INF-γ, IL-6 and IL-1β cytokines levels in the serum of 3xTgAD mice (Fig. 7H). We did not notice any change in the levels of KC/GRO chemokine, and of IL-2, IL-5, and IL-10 cytokines levels in the serum of 3xTgAD mice versus WT mice, and they were all unaffected by Met treatment (Supplementary Fig. 9). To note, the levels of IL-12p70 and of IL-4 cytokines (included in V-plex test) were almost undetectable in the serum of both WT and 3xTgAD mice (data not shown).

Finally, in order to identify potential genes modulated by Met in 3xTgAD hippocampi, we evaluated the gene expression profile using the NanoString mouse inflammatory panel. This panel covers 770 inflammation-associated genes implicated in multiple pathways. We identified 17 statistically differentially expressed genes (*P* <0.05) (Fig. 7I). Among them, 5 genes were downregulated by Met and were shown to be implicated in oligodendrocytes function (OPALIN, ENPP6 and SOX10), DNA damage (RAD51) or yet unknown function (TMEM88B). Met treatment of 3xTgAD mice is associated with the upregulation of genes linked to inflammatory signaling including TYROBP, CTSS, NCF1, AGO4, TREM2,, and CTSS. Interestingly, two of these genes belong to the same signaling cascade (i.e. TYROBP, TREM2) and were described to be dysregulated in AD (26, 27).

These results demonstrate a potential beneficial impact of Met on both central and peripheral inflammation. Other experiments are still needed to determine the specific impact of the identified genes in the regulation of glial cells remodelling due to Met treatment in 3xTgAD brains.

### Metformin ameliorates dendritic spines alterations *ex vivo* and alleviates learning deficit *in vivo*

We studied the functional consequence of Met and CC on synaptic plasticity through the analyses of the morphology of dendritic spines (Figure 1E). To this aim, we used organotypic hippocampal slices cultures (OHSC) that were transduced with lentiviruses expressing Green protein or APPswe protein expressed together with Green protein with IRES system (APPswe-Green) (19) (Fig. 8A and supplementary Fig. 10A). Infected OHSC were then untreated (Control) or treated with Met (Fig. 8A) or with CC (supplementary Fig. 10A). Dendrite imaging allowed us to visualize and index three morphological types of spines reflecting their degree of maturity, i.e. mature spines (M), immature stubby spines (I) or filopodia (F) (see representative dendrites crops in Fig. 8A and supplementary Fig. 10A and the quantification in Fig. 8B and supplementary Fig. 10B). As we already reported (28), we reveal a drastic reduction of the number of mature (M) spines in APPswe-Green dendrites concomitant to a significant increase in the number of filopodia type (F) spines (Fig. 8A, B). The number of immature spines remain unchanged between Green and APPswe-Green dendrites (Fig. 8A, B). Importantly, while we did not report any significant change in spine morphologies between control and Met-treated Green slices, we show in APPswe-Green dendrites that Met treatment significantly increases the number of mature (M) and reduces that of filopodia (F) spines (Fig. 8A, B). CC treatment triggered a significant reduction of mature (M) dendrites and an increase of immature ones in Green dendrites but did not exacerbateed spine alterations in APPswe-Green dendrites (supplementary Fig. 10A, B).

**Fig. 8.**
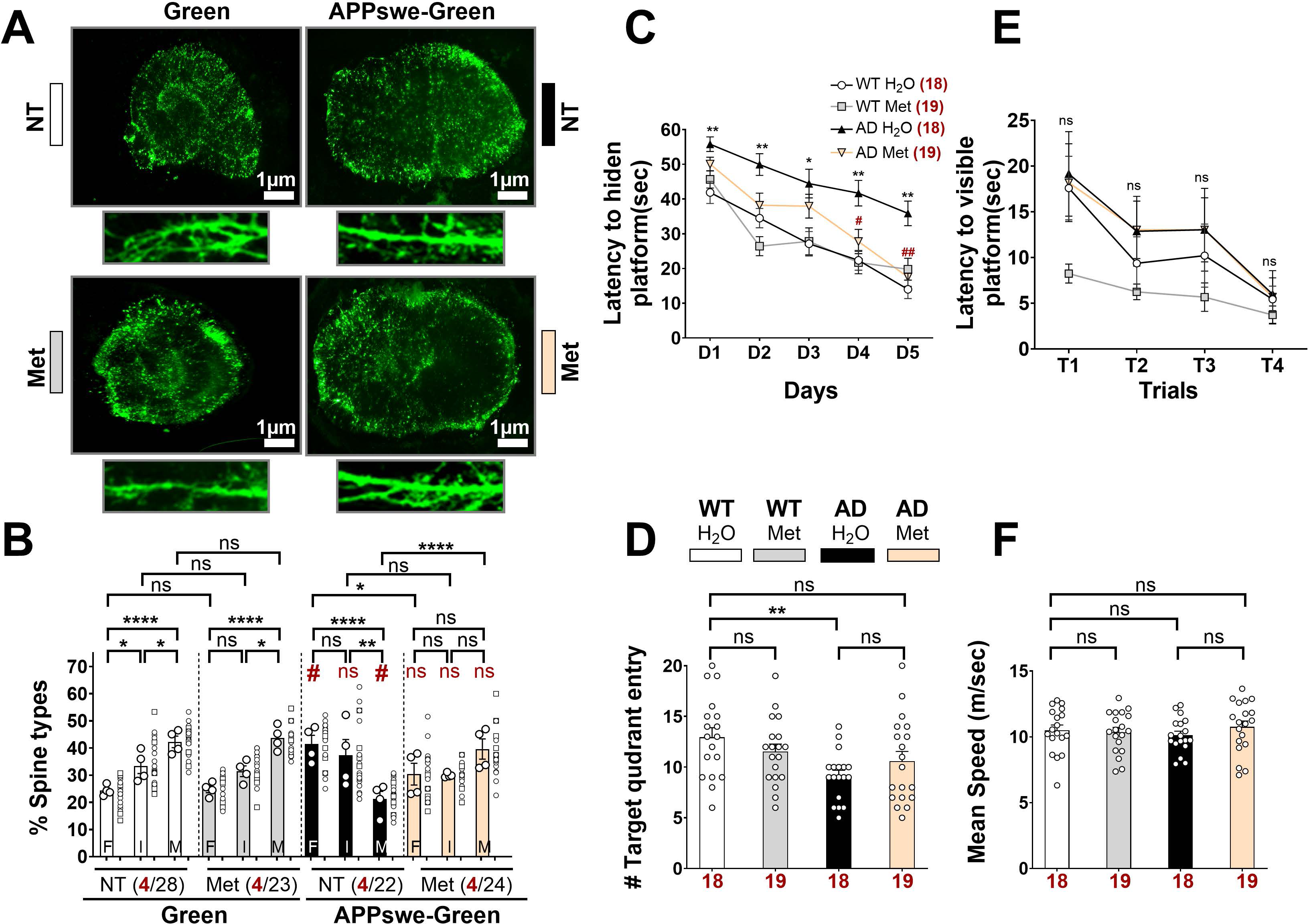
Metformin reverses dendritic spines alterations *ex vivo* and alleviates learning deficit in 3xTgAD mice. (**A**) Representative images of organotypic hippocampal slices (scale bars, 1 µm) and of dendritic segments expressing Green or APPswe-Green mice non-treated (NT) or treated with Met. (2 mM) for 6 h. (**B**) Graph representing the percentage of spine types (F: filopodia, I: immature, and M: mature). (**C**) Latencies to non-visible platform in the MWM test of WT and 3xTgAD (AD) mice untreated (H_2_O) or treated with Met (Fig. 1A). Data are expressed as means of 4 trials per day for 5 consecutive days. (**D**) Number of entries in target quadrant, recorded 24 h after the learning phase. (**E, F**) Latencies to visible platform (E) and swimming speeds (m/sec) (F) recorded 24 h after the learning phase. All data are presented as mean ± SEM. The number of analyzed organotypic slices (B) and mice (C-F) is indicated in red. The number of analyzed dendrites (B) is indicated in black. *P* values were calculated using a 2-Way ANOVA test followed by a Tukey’s multiple comparison test (B), using a mixed model ANOVA with repeated measures (genotype × treatment × training day) and followed by post-hoc comparisons with Holm correction of the *P*-value for multiplicity (C, E), or using Kruskal-Wallis test and Dunn’s multiple comparisons post-test (D, F). * *P* <0.05, ** *P* <0.01, **** *P* <0.0001 and ns: non-significant. (B) **#** *P* <0.05 and ns: non-significant are relative to spine type in the Green NT condition. (C) **#** *P* <0.05 and **##** *P* <0.01 are relative to the AD + H_2_O latencies of the corresponding training day.

The 3xTgAD mice display cognitive alterations in memory and learning tasks (18) that are characteristic defects observed in AD patients. We sought to determine whether Met may positively impact learning and memory using the Morris Water Maze (MWM) as a readout of spatial learning task (18). We report that the average latencies to find the hidden platform were increased in untreated 3xTg-AD and that Met treatment significantly improved the escape latency compared to untreated 3xTgAD mice, while having no effect in WT treated mice (Fig. 8C). We then assessed probe trial 24 h after the last learning day to examine long-term memory and revealed as we already reported (18), that 3xTgAD mice exhibited reduced number of quadrant entries compared to untreated WT mice (Fig. 8D). Although Met treatment failed to normalize the number of target quadrant entry in 3xTgAD mice (Fig. 8D), it abolishes the significant reduction that we observed between 3xTgAD H_2_O and WT H_2_O mice. Of importance, the observed deficits in the MWM in 3xTgAD mice and the impact of Met on learning abilities in 3xTgAD mice were not due to defects in vision (Fig. 8E) or swimming speed (Fig. 8F).

All over, these analyses revealed a beneficial effect of Met on learning ability in 3xTgAD mice, likely attributable to restored synaptic deficits.

## Discussion

Our study unravels altered AMPK-ULK1 cascade in human sporadic AD brains and in mice and cells mimicking familial AD. Our data support previous findings that similarly reported altered AMPK signaling cascade in aging and in AD mice models (29-31). In addition of our demonstration in our human AD brains, anotherstudy have showed that AMPK activation in sporadic AD brains was associated with downstream inhibition of both ULK1 and TBK1 (32). All together, these data converge to suggest a blockade of AMPK and/or of its downstream effector ULK1 in AD.

The activation of the AMP-activated protein kinase (AMPK) is one of the primary mechanism by which Met is suspected to produce beneficial effects (15). A recent study provided a structure-based rationale for the mode of action of Met and its analogues, hence explaining the relative degree of inhibition on complex I (33), causing an increased AMP/ATP and NAD^+^/NADH ratios, and thereby activating AMPK (34). Met has been proven to promote mitochondrial fission, to repair the damaged TCA cycle, to improve mitochondrial respiration, and to restore the mitochondrial life cycle, thereby stabilizing mitochondrial functions (29). We report herein that Met treatment alleviates mitochondria structure alterations in 3xTgAD mice neurons by enhancing mitochondria number, reducing their size and preventing the impairment of cristae morphology. We accordingly demonstrate that Met alleviates mitochondrial membrane potential depolarization in 3xTgAD mice hippocampal neurons. The use of both pharmacological and genetic tools in APPswe cells further supports our results obtained *in vivo* in the 3xTgAD mice.

We further demonstrate that Met prevents the early accumulation of APP-CTFs in the hippocampi of pre-symptomatic 3xTgAD mice aged 4 months and reduces both APP-CTFs and Aβ_1-42_ and Aβ_1-40_ levels in the hippocampi of 3xTgAD mice aged 10-11 months. These results reveal for the first time a beneficial effect of Met on early APP-CTFs accumulation and support previous studies reporting beneficial effects of Met on Aβ pathology in other AD mice models (31, 35, 36). Noticeably, Met-mediated Aβ load decrease was linked to a reduction of BACE-1 expression (36), an increase of insulin degrading enzyme levels (31), or an enhancement of autophagy (37). We postulate herein that the reduction of APP-CTFs load in the hippocampus of pre-symptomatic 3xTgAD mice treated with Met is triggered at least in part by an increase of lysosomal degradation. Indeed, we report a higher localization of APP-CTFs within Cat D^+^ dots and document an increase lysosomes in the hippocampal neuronal soma in Met-treated 3xTgAD mice. We also demonstrate that Met increases Cat D activity *in vitro*. A recent study proposed that Met at low concentration can bind to PEN2 protein in the lysosomal compartment, form a complex with ATP6AP1 (a subunit of the lysosomal v-ATPase), thus inhibiting lysosomal acidification and the activation of AMPK without affecting AMP cellular levels (38). Other studies demonstrate that AMPK can also be activated at the lysosome, where it interacts with other proteins, such as AXIN and LKB1 (39). AMPK-linked autophagy/lysosomes function was also shown to occur through the transcription factor EB (TFEB), known to regulate the expression of several lysosomal proteins, including Cat D (40, 41).

The results of several studies are in favor of AMPK stimulation as a way to reduce Tau pathology in AD. We report that Met selectively reduced pSer202/pThr205-Tau staining in hippocampal neurons *in vivo*. Similarly, Met treatment in 9-month-old APP/PS1 mice, injected with aggregated Tau in the hippocampus, reduces pTau (Ser202/Thr205, Thr231 and Ser422) levels through an enhancement of microglial phagocytosis and clearance of pTau (37). In addition, STZ-mice overexpressing AMPK also showed reduced pThr231- and pSer396-Tau levels (30). A recent study confirmed that Met reduces the hyperphosphorylation of Tau at Ser202/Thr205, Thr231, and Ser422 sites in the soluble fraction and of Ser202/Thr205, Ser262, Thr396, Thr231, and Ser422 sites in the insoluble fraction of tau-seeded PS19 mice brains (42). *In vitro* studies further support these results and show that the treatment of human cortical stem cells with another AMPK activator (AICAR) reduces the phosphorylation levels of Tau on Thr231, Ser199/202, and Ser396 (43).

It should be noted that some studies reported that high concentration of Met increases amyloidogenic APP processing through enhanced APP, BACE-1, and PS1 (γ-secretase) levels *in vitro* (44), and that chronic administration of Met promotes amyloidogenic APP processing in autophagosomes in AD mice at advanced disease stages (45). These discrepancies point-out the importance of a careful Met dosing and treatment time alongside with AD progression and may suggest a differential impact of Met under non-pathological conditions versus AD-related conditions warranting further studies.

In AD, the amyloid pathology is associated with deleterious chronic neuroinflammation (13). We report here that Met reduces hippocampal microgliosis and astrogliosis as well as Aβ plaques load in 3xTgAD mice, likely through the stimulation of the microglial clearance. A similar result has been reported in rats injected with streptozotocin (STZ) modeling SAD (46, 47). Moreover, we show a region-specific modulation of Iba1^+^ cells morphology by Met, reducing the size of ameboid nuclei in the subiculum area near Aβ plaques and the hyper-ramification processes in the DG and CA1 region. The molecular signature and phagocytic activity of Aβ plaques-associated microglia has been extensively studied in AD and are now referred as Disease-Associated microglia (DAM), that undergo protective role early in Alzheimer’s, but contribute to late-stage chronic inflammation thought to accelerate neurodegeneration (48). The functional role of hyper-ramified microglial cells that we describe in DG and CA1 regions remain to be studied in the context of AD (49). The mechanism underlying the anti-inflammatory action of AMPK includes the indirect inhibition of NF-κB, through multiple downstream pathways including SIRT1, Folkhead box O (FOXO) and PGC1α (50). While we did not identify a differential expression of these genes between Met-treated and untreated 3xTgAD mice, we unravel an upregulation of a subset of genes including TREM2 and its adaptor TYROBP. Recent studies converge to show a critical role of TREM2 in AD development (26). In addition, TYROBP interacts closely with receptors involved in phagocytosis, and a substantial body of evidence indicates that several key receptors involved in amyloid clearance or compaction are closely linked to TYROBP (27, 51). In support to the anti-inflammatory potential of Met, we also show that the levels of peripheral pro-inflammatory cytokines (TNF-α, IFN-γ, IL1-β et IL-6) are reduced in 3xTgAD treated mice. The molecular mechanisms behind these regulations are under investigation.

Synaptic plasticity alterations and loss of synapses are major features of AD, that correlate with the cognitive impairments (52, 53). Met reduces the proportion of filopodia dendritic spines *ex vivo* and improves the spatial learning capacities of symptomatic 3xTgAD mice. Accordingly, other drugs activating AMPK (AICAR and resveratrol), analogously dampen spatial memory defects in STZ-rats and CB2R^-/-^ mice (54).

## Conclusion

Our results support a beneficial impact of targeting AMPK-ULK1 cascade at both pre-symptomatic and symptomatic AD stages that extends from bioenergetic, proteinopathy and inflammation alterations to cognitive deficits. Met has garnered interest for its potential to treat neurodegenerative diseases in humans (55). However, while the majority of clinical studies suggests a beneficial effect of Met on the prevention of AD, including improved cognitive performance and decreased AD risks (56-59), some studies reported that long-term use of Met may be linked to a higher AD risk (60). This highlights the need for further research on the repurposing of alternative drugs targeting AMPK-ULK1 cascade as a strategy to reverse or at least to slow down the course of AD onset and/or development in preclinical study models and at early AD stages in humans.

## Funding

This work was supported by Inserm (MC, and FC), Fondation Vaincre Alzheimer (grant # FR18-035) (MC), LABEX (excellence laboratory, program investment for the future) DISTALZ (Development of Innovative Strategies for a Transdisciplinary approach to Alzheimer’s disease) (FC and LB). We acknowledge PhD fellow support from DISTALZ to AM (MC and VBS). We acknowledge the University’s CCMA, Electron Microscopy facility supported by Université Côte d’Azur, the “Conseil Departmental 06”, and GIS IBiSA, and we thank Christelle Boscagli for technical help.

## Author Contributions

Conceptualization: MC; Methodology: AM, MC; Investigation: AM, SB, AV, CB, SLG, SE; Visualization: AM, SB, SLG; Funding acquisition: MC, FC, LB; Supervision: MC; Writing – original draft: AM; Writing – review & editing: MC. Editing: FC, LB, VBS.

## Competing interests

Authors declare no conflict of interest

## Compliance with ethical standards

We follow applicable international, national, and institutional guidelines for the care and use of animals. Experimental procedures performed in studies involving animals were approved by Nice university Animal care and use Committee, and the National Council on animal care of the Ministry of Health (Project n°: APAFIS#20495-201904231352370) in accordance with the guidelines established by the European community council (Directive of November 24^th^, 1986) and with the ethical standards and animal welfare committee of our institute.

All procedures performed in studies involving human participants were in accordance with the ethical standards of the institutional and national research committee and with the 1964 Helsinki declaration and its later amendments or comparable ethical studies. Brain samples were obtained from the Brain Bank “Neuro-CEB” run by a consortium of Patients Associations: ARSEP (association for research on multiple sclerosis), CSC (cerebellar ataxias), and France Parkinson. The consents were signed by the patients themselves or their next of kin in their name, in accordance with the French Bioethical (agreement AC-2013-1887). Cases were anonymized but information was provided regarding sex, age at death, and neuropathology (Suppl.Table 1). Informed consent for tissue donation for research is obtained by the brain banks under their approval procedures.

## Statement

We Declare that we can provide original data and supporting material that can be accessible.

## Notes

### Competing Interest Statement

The authors have declared no competing interest.

## References

1. Scheltens P, De Strooper B, Kivipelto M, Holstege H, Chetelat G, Teunissen CE, et al. Alzheimer’s disease. Lancet. 2021;397(10284):1577–90.

2. Bloom GS. Amyloid-beta and tau: the trigger and bullet in Alzheimer disease pathogenesis. JAMA Neurol. 2014;71(4):505–8.

3. Zhang YW, Thompson R, Zhang H, Xu H. APP processing in Alzheimer’s disease. Mol Brain. 2011;4:3.

4. Pulina MV, Hopkins M, Haroutunian V, Greengard P, Bustos V. C99 selectively accumulates in vulnerable neurons in Alzheimer’s disease. Alzheimers Dement. 2020;16(2):273–82.

5. Lauritzen I, Pardossi-Piquard R, Bauer C, Brigham E, Abraham JD, Ranaldi S, et al. The beta-secretase-derived C-terminal fragment of betaAPP, C99, but not Abeta, is a key contributor to early intraneuronal lesions in triple-transgenic mouse hippocampus. J Neurosci. 2012;32(46):16243-1655a.

6. Vaillant-Beuchot L, Mary A, Pardossi-Piquard R, Bourgeois A, Lauritzen I, Eysert F, et al. Accumulation of amyloid precursor protein C-terminal fragments triggers mitochondrial structure, function, and mitophagy defects in Alzheimer’s disease models and human brains. Acta Neuropathol. 2021;141(1):39–65.

7. Kwart D, Gregg A, Scheckel C, Murphy EA, Paquet D, Duffield M, et al. A Large Panel of Isogenic APP and PSEN1 Mutant Human iPSC Neurons Reveals Shared Endosomal Abnormalities Mediated by APP beta-CTFs, Not Abeta. Neuron. 2019;104(5):1022.

8. Bourgeois A, Lauritzen I, Lorivel T, Bauer C, Checler F, Pardossi-Piquard R. Intraneuronal accumulation of C99 contributes to synaptic alterations, apathy-like behavior, and spatial learning deficits in 3xTgAD and 2xTgAD mice. Neurobiol Aging. 2018;71:21–31.

9. Kaur G, Pawlik M, Gandy SE, Ehrlich ME, Smiley JF, Levy E. Lysosomal dysfunction in the brain of a mouse model with intraneuronal accumulation of carboxyl terminal fragments of the amyloid precursor protein. Mol Psychiatry. 2017;22(7):981–9.

10. Mondragon-Rodriguez S, Gu N, Manseau F, Williams S. Alzheimer’s Transgenic Model Is Characterized by Very Early Brain Network Alterations and beta-CTF Fragment Accumulation: Reversal by beta-Secretase Inhibition. Front Cell Neurosci. 2018;12:121.

11. Lauritzen I, Pardossi-Piquard R, Bourgeois A, Pagnotta S, Biferi M-G, Barkats M, et al. Intraneuronal aggregation of the β-CTF fragment of APP (C99) induces Aβ-independent lysosomal-autophagic pathology. Acta Neuropathologica. 2016;132:257–76.

12. Wang W, Zhao F, Ma X, Perry G, Zhu X. Mitochondria dysfunction in the pathogenesis of Alzheimer’s disease: recent advances. Molecular Neurodegeneration. 2020;15(1):30.

13. Bettcher BM, Tansey MG, Dorothee G, Heneka MT. Peripheral and central immune system crosstalk in Alzheimer disease - a research prospectus. Nat Rev Neurol. 2021.

14. Trefts E, Shaw RJ. AMPK: restoring metabolic homeostasis over space and time. Mol Cell. 2021;81(18):3677–90.

15. Rotermund C, Machetanz G, Fitzgerald JC. The Therapeutic Potential of Metformin in Neurodegenerative Diseases. Front Endocrinol (Lausanne). 2018;9:400.

16. Oddo S, Caccamo A, Shepherd JD, Murphy MP, Golde TE, Kayed R, et al. Triple-transgenic model of Alzheimer’s disease with plaques and tangles: intracellular Abeta and synaptic dysfunction. Neuron. 2003;39(3):409–21.

17. Labuzek K, Suchy D, Gabryel B, Bielecka A, Liber S, Okopien B. Quantification of metformin by the HPLC method in brain regions, cerebrospinal fluid and plasma of rats treated with lipopolysaccharide. Pharmacol Rep. 2010;62(5):956–65.

18. Lacampagne A, Liu X, Reiken S, Bussiere R, Meli AC, Lauritzen I, et al. Post-translational remodeling of ryanodine receptor induces calcium leak leading to Alzheimer’s disease-like pathologies and cognitive deficits. Acta Neuropathol. 2017;134(5):749–67.

19. Bussiere R, Oules B, Mary A, Vaillant-Beuchot L, Martin C, El Manaa W, et al. Upregulation of the Sarco-Endoplasmic Reticulum Calcium ATPase 1 Truncated Isoform Plays a Pathogenic Role in Alzheimer’s Disease. Cells. 2019;8(12).

20. Schatzle P, Kapitein LC, Hoogenraad CC. Live imaging of microtubule dynamics in organotypic hippocampal slice cultures. Methods Cell Biol. 2016;131:107–26.

21. Pinto B, Morelli G, Rastogi M, Savardi A, Fumagalli A, Petretto A, et al. Rescuing Over-activated Microglia Restores Cognitive Performance in Juvenile Animals of the Dp(16) Mouse Model of Down Syndrome. Neuron. 2020;108(5):887–904 e12.

22. Oules B, Del Prete D, Greco B, Zhang X, Lauritzen I, Sevalle J, et al. Ryanodine receptor blockade reduces amyloid-beta load and memory impairments in Tg2576 mouse model of Alzheimer disease. J Neurosci. 2012;32(34):11820–34.

23. Gomez-Murcia V, Launay A, Carvalho K, Burgard A, Meriaux C, Caillierez R, et al. Neuronal A2A receptor exacerbates synapse loss and memory deficits in APP/PS1 mice. Brain : a journal of neurology. 2024;147(8):2691–705.

24. Sepulveda-Diaz JE, Ouidja MO, Socias SB, Hamadat S, Guerreiro S, Raisman-Vozari R, et al. A simplified approach for efficient isolation of functional microglial cells: Application for modeling neuroinflammatory responses in vitro. Glia. 2016;64(11):1912–24.

25. Grubman A, Choo XY, Chew G, Ouyang JF, Sun G, Croft NP, et al. Transcriptional signature in microglia associated with Abeta plaque phagocytosis. Nat Commun. 2021;12(1):3015.

26. Griciuc A, Patel S, Federico AN, Choi SH, Innes BJ, Oram MK, et al. TREM2 Acts Downstream of CD33 in Modulating Microglial Pathology in Alzheimer’s Disease. Neuron. 2019;103(5):820–35 e7.

27. Haure-Mirande JV, Audrain M, Ehrlich ME, Gandy S. Microglial TYROBP/DAP12 in Alzheimer’s disease: Transduction of physiological and pathological signals across TREM2. Mol Neurodegener. 2022;17(1):55.

28. Valverde A, Dunys J, Lorivel T, Debayle D, Gay AS, Lacas-Gervais S, et al. Aminopeptidase A contributes to biochemical, anatomical and cognitive defects in Alzheimer’s disease (AD) mouse model and is increased at early stage in sporadic AD brain. Acta Neuropathol. 2021.

29. Wang Y, An H, Liu T, Qin C, Sesaki H, Guo S, et al. Metformin Improves Mitochondrial Respiratory Activity through Activation of AMPK. Cell Rep. 2019;29(6):1511–23 e5.

30. Wang L, Li N, Shi FX, Xu WQ, Cao Y, Lei Y, et al. Upregulation of AMPK Ameliorates Alzheimer’s Disease-Like Tau Pathology and Memory Impairment. Mol Neurobiol. 2020;57(8):3349–61.

31. Lu XY, Huang S, Chen QB, Zhang D, Li W, Ao R, et al. Metformin Ameliorates Aβ Pathology by Insulin-Degrading Enzyme in a Transgenic Mouse Model of Alzheimer’s Disease. Oxid Med Cell Longev. 2020;2020:2315106.

32. Fang EF, Hou Y, Palikaras K, Adriaanse BA, Kerr JS, Yang B, et al. Mitophagy inhibits amyloid-β and tau pathology and reverses cognitive deficits in models of Alzheimer’s disease. Nature Neuroscience. 2019;22(3):401–12.

33. Bridges HR, Blaza JN, Yin Z, Chung I, Pollak MN, Hirst J. Structural basis of mammalian respiratory complex I inhibition by medicinal biguanides. Science (New York, N Y). 2023;379(6630):351-7.

34. Foretz M, Guigas B, Bertrand L, Pollak M, Viollet B. Metformin: from mechanisms of action to therapies. Cell Metab. 2014;20(6):953–66.

35. Chen JL, Luo C, Pu D, Zhang GQ, Zhao YX, Sun Y, et al. Metformin attenuates diabetes-induced tau hyperphosphorylation in vitro and in vivo by enhancing autophagic clearance. Exp Neurol. 2019;311:44–56.

36. Ou Z, Kong X, Sun X, He X, Zhang L, Gong Z, et al. Metformin treatment prevents amyloid plaque deposition and memory impairment in APP/PS1 mice. Brain Behav Immun. 2018;69:351–63.

37. Chen Y, Zhao S, Fan Z, Li Z, Zhu Y, Shen T, et al. Metformin attenuates plaque-associated tau pathology and reduces amyloid-β burden in APP/PS1 mice. Alzheimers Res Ther. 2021;13(1):40.

38. Ma T, Tian X, Zhang B, Li M, Wang Y, Yang C, et al. Low-dose metformin targets the lysosomal AMPK pathway through PEN2. Nature. 2022;603(7899):159-65.

39. Zhang YL, Guo H, Zhang CS, Lin SY, Yin Z, Peng Y, et al. AMP as a low-energy charge signal autonomously initiates assembly of AXIN-AMPK-LKB1 complex for AMPK activation. Cell Metab. 2013;18(4):546–55.

40. Malik N, Ferreira BI, Hollstein PE, Curtis SD, Trefts E, Weiser Novak S, et al. Induction of lysosomal and mitochondrial biogenesis by AMPK phosphorylation of FNIP1. Science. 2023;380(6642):eabj5559.

41. Paquette M, El-Houjeiri L, L CZ, Puustinen P, Blanchette P, Jeong H, et al. AMPK-dependent phosphorylation is required for transcriptional activation of TFEB and TFE3. Autophagy. 2021;17(12):3957–75.

42. Zhao S, Fan Z, Zhang X, Li Z, Shen T, Li K, et al. Metformin Attenuates Tau Pathology in Tau-Seeded PS19 Mice. Neurotherapeutics. 2023;20(2):452–63.

43. Kim B, Figueroa-Romero C, Pacut C, Backus C, Feldman EL. Insulin Resistance Prevents AMPK-induced Tau Dephosphorylation through Akt-mediated Increase in AMPKSer-485 Phosphorylation. J Biol Chem. 2015;290(31):19146–57.

44. Picone P, Nuzzo D, Caruana L, Messina E, Barera A, Vasto S, et al. Metformin increases APP expression and processing via oxidative stress, mitochondrial dysfunction and NF-κB activation: Use of insulin to attenuate metformin’s effect. Biochim Biophys Acta. 2015;1853(5):1046–59.

45. Son SM, Shin HJ, Byun J, Kook SY, Moon M, Chang YJ, et al. Metformin Facilitates Amyloid-β Generation by β- and γ-Secretases via Autophagy Activation. J Alzheimers Dis. 2016;51(4):1197–208.

46. Saffari PM, Alijanpour S, Takzaree N, Sahebgharani M, Etemad-Moghadam S, Noorbakhsh F, et al. Metformin loaded phosphatidylserine nanoliposomes improve memory deficit and reduce neuroinflammation in streptozotocin-induced Alzheimer’s disease model. Life Sci. 2020;255:117861.

47. Pilipenko V, Narbute K, Pupure J, Langrate IK, Muceniece R, Kluša V. Neuroprotective potential of antihyperglycemic drug metformin in streptozocin-induced rat model of sporadic Alzheimer’s disease. Eur J Pharmacol. 2020;881:173290.

48. Jay TR, Hirsch AM, Broihier ML, Miller CM, Neilson LE, Ransohoff RM, et al. Disease Progression-Dependent Effects of TREM2 Deficiency in a Mouse Model of Alzheimer’s Disease. J Neurosci. 2017;37(3):637–47.

49. Fujikawa R, Jinno S. Identification of hyper-ramified microglia in the CA1 region of the mouse hippocampus potentially associated with stress resilience. Eur J Neurosci. 2022;56(8):5137–53.

50. Peixoto CA, Oliveira WH, Araujo S, Nunes AKS. AMPK activation: Role in the signaling pathways of neuroinflammation and neurodegeneration. Exp Neurol. 2017;298(Pt A):31–41.

51. Audrain M, Haure-Mirande JV, Mleczko J, Wang M, Griffin JK, St George-Hyslop PH, et al. Reactive or transgenic increase in microglial TYROBP reveals a TREM2-independent TYROBP-APOE link in wild-type and Alzheimer’s-related mice. Alzheimers Dement. 2021;17(2):149–63.

52. Onyango IG, Jauregui GV, Carna M, Bennett JP, Jr., Stokin GB. Neuroinflammation in Alzheimer’s Disease. Biomedicines. 2021;9(5).

53. Kinney JW, Bemiller SM, Murtishaw AS, Leisgang AM, Salazar AM, Lamb BT. Inflammation as a central mechanism in Alzheimer’s disease. Alzheimers Dement (N Y). 2018;4:575–90.

54. Wang H, Jiang T, Li W, Gao N, Zhang T. Resveratrol attenuates oxidative damage through activating mitophagy in an in vitro model of Alzheimer’s disease. Toxicology Letters. 2018;282:100–8.

55. Du MR, Gao QY, Liu CL, Bai LY, Li T, Wei FL. Exploring the Pharmacological Potential of Metformin for Neurodegenerative Diseases. Front Aging Neurosci. 2022;14:838173.

56. Zhou JB, Tang X, Han M, Yang J, Simo R. Impact of antidiabetic agents on dementia risk: A Bayesian network meta-analysis. Metabolism. 2020;109:154265.

57. Kuan YC, Huang KW, Lin CL, Hu CJ, Kao CH. Effects of metformin exposure on neurodegenerative diseases in elderly patients with type 2 diabetes mellitus. Prog Neuropsychopharmacol Biol Psychiatry. 2017;79(Pt B):77–83.

58. Koenig AM, Mechanic-Hamilton D, Xie SX, Combs MF, Cappola AR, Xie L, et al. Effects of the Insulin Sensitizer Metformin in Alzheimer Disease: Pilot Data From a Randomized Placebo-controlled Crossover Study. Alzheimer Dis Assoc Disord. 2017;31(2):107–13.

59. Hsu CC, Wahlqvist ML, Lee MS, Tsai HN. Incidence of dementia is increased in type 2 diabetes and reduced by the use of sulfonylureas and metformin. J Alzheimers Dis. 2011;24(3):485–93.

60. Imfeld P, Bodmer M, Jick SS, Meier CR. Metformin, other antidiabetic drugs, and risk of Alzheimer’s disease: a population-based case-control study. J Am Geriatr Soc. 2012;60(5):916–21.

