## Supplementary material and figures for "Hampered AMPK-ULK1 cascade in Alzheimer’s disease (AD) instigates mitochondria dysfunctions and AD-related alterations that are alleviated by metformin"

**List of Supplementary Materials**

**Supplementary methods**

**Supplementary table 1.** Demographic data and neuropathology evolution of human brain samples

**Supplementary Table 2.** List of antibodies used in the study.

**Supplementary Fig. 1.** Met treatment does not impact the daily drink consumption and bodyweight evolution in mice.

**Supplementary Fig. 2.** AMPK-ULK1 cascade in altered in SH-SY5Y cells expressing APPswe and effect of Met and CC.

**Supplementary Fig. 3.** AMPK-ULK1 cascade modulation impacts mitochondrial structure and function in APPswe expressing cells.

**Supplementary Fig. 4.** CC triggers mitochondrial function alteration in control cells.

**Supplementary Fig. 5.** Genetic modulation of AMPK-ULK1 cascade impacts mitochondrial function in APPswe expressing cells.

**Supplementary Fig. 6.** Met does impact the levels of Tau phosphorylation on Ser199, Thr231, and Ser396 residues in the hippocampi of 3xTgAD mice.

**Supplementary Fig. 7.** Metn alleviates astrogliosis in 3xTgAD mice.

**Supplementary Fig. 8.** CC reduces phagocytic activity of microglia.

**Supplementary Fig. 9.** Met treatment does not impact the circulating levels of IL-10, KC/GRO, IL-2, and IL-5.

**Supplementary Fig. 10.** CC triggers spine morphology alterations in *ex vivo* organotypic hippocampal slice cultures.

#### Supplementary methods

##### Cell lines and treatments

Human SH-SY5Y cells stably expressing pcDNA3.1 (Control) or full-length APPswe constructs were generated, and maintained in the presence of 400 µg/mL geneticin (MP Biomedicals as already described) (1). Cells were treated for 6 h with Dorsomorphin known also as Compound C (CC) (Abcam) at 10 µM, or with metformin-HCl (Met) (Merck-Millipore) at 2 mM.

##### Viral production of AMPK constructs

we produce viruses expressing wild type AMPK$\alpha_2$ (WT-AMPK), AMPK$\alpha_2$ constitutively active form (CA-AMPK) obtained by the truncation after aa 312 residue and dominant negative K45R AMPK$\alpha_2$ mutant (AMPK-DN) cloned in SIN-PGK-cPPT-RFA-WHV lentiviral vector, under the control of the PGK promoter (mutants were sub-cloned from Addgene plasmid 159991-Morris J. Birnbaum (2)). Viral titers were similarly assessed using p24 ELISA (Cell Biolabs, San Diego, Cat#VPK-107).

##### Flux cytometry measurements of mitochondrial superoxide and of mitochondrial potential in cells

MitoSOX Red is a highly selective dye detecting superoxide in the mitochondria of living cells (3). Cells were incubated in 5 µM MitoSOX red mitochondrial superoxide indicator (Invitogen) in DMEM for 30 min, at37°C, 5% $CO_2$. Cells were harvested and rinsed twice with ice cold HBSS complemented with 1 mM $CaCl_2$ and 0.5 mM $MgCl_2$. For TMRM analyses, cells were harvested, rinsed with PBS, and incubated in TMRM (prepared in DMEM) (2 nM) for 30 min at 37°C, 5% $CO_2$. TMRM and MitoSOX fluorescence median intensities were then analyzed in the Novocyte Flow Cytometer (ACEA bioscience,Inc) excitation/emission 510 nm/580 nm.

To ascertain for the specific TMRM and MitoSOX signals, cells were respectively treated with the mitochondrial uncoupler FCCP (trifluoromethoxy carbonylcyanide phenylhydrazone), or with Oligomycin A (ATPase inhibitor) and antimycin (inhibitor of cytochrome C reductase).

**Analyses of mitochondria ultrastructure by transmission electron microscopy in Cells**

We follow the protocol for cell processing previously described (3).

**Supplementary table 1.** Demographic data and neuropathology evolution of human brain samples described by NFT Braak's stage, Thal Aβ phase, and Amyloid angiopathy. Controls are brain samples isolated from post-mortem patients diagnosed as negative for AD pathology; (*) suspected to be a familial AD case. NA: not available; PMD: Post-mortem delay in hours (h). We used region of the temporal lobe (T1).

| | Gender | Age (years) | PMD (h) | **Brain region** | **NFT Braak's stage** | **Thal Aβ phase** | **Amyloid angiopathy** |
|---|---|---|---|---|---|---|---|
| **Control** | Male | 71 | 26 | T1 | - | - | NA |
| | Male | 84 | 32 | T1 | - | - | NA |
| | Female | 71 | 15 | T1 | - | 3 | 2 |
| | Female | 60 | 28 | T1 | - | 0 | 0 |
| | Female | 89 | 12 | T1 | - | NA | NA |
| | Female | 62 | 44 | T1 | - | 0 | 0 |
| | Male | 55 | 21 | T1 | - | 1 | 0 |
| **AD** | Female | 89 | 26 | T1 | Stade VI | NA | NA |
| | Female | 81 | 60 | T1 | Stade IV | NA | NA |
| | Female | 80 | 51 | T1 | Stade V | NA | NA |
| | Female | 84 | 81 | T1 | Stade IV | 4 | 2 |
| | Female | 65 | 41 | T1 | Stade VI | 5 | 1 |
| | Male | 81 | 19 | T1 | Stade VI | NA | NA |
| | Female | 75 | 7 | T1 | Stade VI | 5 | 2 |
| | Female | 93 | 21 | T1 | Stade VI | NA | NA |
| | Female | 91 | 34 | T1 | Stade VI | 5 | 1 |
| | Female | 55* | 58 | T1 | Stade VI | 5 | 1 |
| | Female | 81 | NA | T1 | Stade VI | 5 | 1 |
| | Female | 82 | NA | T1 | Stade VI | 4 | 1 |
| | Male | 86 | 32 | T1 | Stade VI | 5 | 1 |

**Supplementary Table 2:** List of antibodies used in the study.

| ANTIBODY | SOURCE | IDENTIFIER | DILUTION |
|---|---|---|---|
| Rabbit anti-phospho T172 AMPKα | Cell Signaling Technology | Cat#50081S | 1:1000 (WB) |
| Rabbit anti-AMPKα | Cell Signaling Technology | Cat#2532 | 1:1000 (WB) |
| Rabbit anti-phospho S555 ULK1 (D1H4) | Cell Signaling Technology | Cat#5869 | 1:1000 (WB) |
| Rabbit anti-ULK1 (D8H5) | Cell Signaling Technology | Cat#8054 | 1:1000 (WB) |
| Rabbit anti-GFAP | Novus Biologicals | Cat#NB300-141 | 1:1000 (IHC) |
| Mouse monoclonal anti-pTau (Ser202/Thr205) (AT8) | Thermo Fisher Scientific | Cat#MN1020 | 1:1000 (IHC) |
| Rabbit Recombinant Monoclonal Cathepsin D antibody | Abcam | EPR3057Y | 1/1000 (IF) |
| Mouse Monoclonal anti-β-Actin | Sigma-Aldrich | Cat#A5316 | 1:5000 (WB) |
| Goat HRP-Conjugated anti-Rabbit IgG | Jackson ImmunoResearch | Cat#111-036-045; RRID: AB_2337943 | 1:1000 (IHC) |
| Goat HRP-Conjugated anti-Mouse IgG | Jackson ImmunoResearch | Cat#115-036-003; RRID: AB_2338518 | 1:1000 (IHC) |

**Legends to Supplementary figures**

**Supplementary Fig. 1. Met treatment does not impact the daily drink consumption and bodyweight evolution in mice**. (**A**) Evolution of the volume of the water consumption per mouse per day during one week. All data are presented as mean ± SEM. The number of mice is indicated in red. (**B**) Mice bodyweight evolution during the 4 weeks of Met treatment in mice aged 10 to 11 months.

**Supplementary Fig. 2. AMPK-ULK1 cascade in altered in SH-SY5Y cells expressing APPswe and effect of Metformin and CC.** (**A**) Representative SDS-PAGE showing full-length APP, APP-CTFs (i.e. C99 and C83), p(Thr172)AMPK, total AMPK, p(Ser555)ULK1, and total ULK1 protein levels. Actin was used as loading control. (**B, C**) Quantitative graphs of indicated proteins and the ratio of the phosphorylated forms of AMPK, ULK1 versus respective total proteins in control (n=8) and APPswe expressing cells (n=8). (**D, G**) Representative SDS-PAGE p(Thr172)AMPK, total AMPK, p(Ser555)ULK1, and total ULK1 protein levels in APPswe expressing cells treated with 2 mM Met (D) or 10 µM CC (G) for 6 h. Actin was used as loading control. (**E, F, H, I**) Quantitative graphs of indicated proteins as in B, C) in control (n=6-7) and APPswe expressing cells treated with Met or CC (n=6-7). Graphs represent means ± S.E.M versus control cells (taken as 1). * $P$ <0.05, ** $P$ <0.01, **** $P$ <0.0001 and ns: non-significant using Mann Whitney test.

**Supplementary Fig. 3. AMPK-ULK1 cascade modulation impacts mitochondrial structure and function in APPswe expressing cells.** (**A, D**) Electron microscopy ultrastructure of APPswe expressing cells treated for 6h with vehicle or CC (10 µM) (A), or untreated and treated with Met (2 mM) (D). Scale bars correspond to2 µm. N: nucleus. Yellow and white arrowheads indicate mitochondria (white: mitochondria with smaller area versus yellow ones). (**B, C, E, F**) Quantitative graphs of the mitochondria number/10 $\mu m^2$ (B, E) and area (µm$^2$) (C, F) in APPswe expressing cells treated as in (A), or as in (D). Graphs represents means ± SEM obtained from 2 independent experiments (indicated in red) and different images (B, E) or isolated mitochondria (C, F) (black) (**G, H**) Graphs representing TMRM (G) or MitoSOX (H) median intensities obtained by FACS analyses in cells expressing empty vector (control) and in APPswe expressing cells untreated or treated for 6h with vehicle, CC (10 µM) or Met (2 µm). Median fluorescence intensities are expressed as means ± SEM versus control (taken as 1) obtained from at least three independent experiments and independent wells (indicated in red). (B, C, E, G, H) * *P* <0.05, ** *P* <0.01, *** *P* <0.001, **** *P* <0.0001 and ns: non-significant using Mann Whitney test or krustal-Wallis and Dunn's multiple comparisons post-test.

**Supplementary Fig. 4. CC triggers mitochondrial function alteration in control cells.** (**A, B**) Graphs representing TMRM (A) or MitoSOX (B) median intensities obtained by FACS analyses in cells expressing empty vector (control) untreated or treated for 6h with vehicle, CC (10 µM) or Met (2 mM). Median fluorescence intensities are expressed as means ± SEM versus control (taken as 1) obtained from at least three independent experiments and independent wells (indicated in red). *** *P* <0.001, and ns: non-significant using Mann Whitney test.

**Supplementary Fig. 5. Genetic modulation of AMPK-ULK1 cascade impacts mitochondrial function in APPswe expressing cells.** (**A**) Representative SDS-PAGE showing the overexpression of WT-AMPK, truncated CA-AMPK and DN-AMPK constructs. (**B, C**) Quantitative graphs of p(Thr172)AMPK, p(Ser555)ULK1, and total ULK1, protein levels and the ratio of the phosphorylated forms of AMPK, and ULK1 versus respective total proteins. Actin is used as loading control. (**D, E**) Graphs representing TMRM (D) or MitoSOX (E) median intensities obtained by FACS analyses in cells transfected as in (A). Median fluorescence intensities are expressed as means ± SEM versus control (taken as 1). Data are obtained from independent wells (indicated in red). * $P < 0.05$, ** $P < 0.01$ and ns: non-significant using Kruskal-Wallis test and Dunn's multiple comparisons post-test.

**Supplementary Fig. 6. Met does impact the levels of Tau phosphorylation on Ser199, Thr231, and Ser396 residues in the hippocampi of 3xTgAD mice.** (**A**, **B**) Quantitative graphs of the levels of phosphorylated Tau species (Ser199, Thr231, Ser396) in the hippocampi homogenates of AD + $H_2O$ (n = 14) and AD + Met (n = 14), normalized by the total (murine and human isoforms) Tau level (A), or by the human Tau level (B). All data are presented as mean ± SEM. *P* values were calculated using a Mann Whitney test. ns: non-significant. The number analyzed mice is indicated in red.

**Supplementary Fig. 7. Met alleviates astrogliosis in 3xTgAD mice.** (**A**) Representative images showing GFAP in the hippocampus and higher magnification of the subiculum area of 10-11-month-old WT and 3xTgAD mice untreated ($H_2O$) or treated with Met. Scale bars, 500 µm (top) or 50 µm (bottom). (**B**) Quantitative graph of GFAP positive cells / $mm^2$ of subiculum. Data are presented as mean ± SEM. The number of analyzed mice is indicated in bold, and the number of slices is in light. *P* values were calculated using a 2-Way ANOVA test followed by a Tukey's multiple comparison test. *** $P < 0.001$, **** $P < 0.0001$, and ns: non-significant. The number of analyzed mice is indicated in red. The number of analyzed slices is in black.

**Supplementary Fig. 8. CC reduces phagocytic activity of microglia.** (**A**) Representative images of primary microglia incubated with fluorescent microbeads (Green), treated or not with 10 µM CC for 6 h and stained with Iba1 antibody (Red). Nuclei were stained with DAPI. Scale bars, 10 µm. (**F**) Quantitative graph of the proportion of phagocytic microglia (that contains at least one microbead). (**G**) Quantitative graph of the number of microbeads per microglia. *P* values were calculated using a Mann Whitney test **** $P < 0.0001$.

**Supplementary Fig. 9. Met treatment does not impact the circulating levels of IL-10, KC/GRO, IL-2, and IL-5.** Graph Plots representing the levels of inflammatory markers (IL-10, KC/GRO, IL-2, and IL-5) (pg/ml) in the serum of WT and 3xTgAD mice untreated ($H_2O$) or treated with Met. The number of analyzed mice is indicated in Fig. 5H. *P* values were calculated using a Mann Whitney multiple comparisons post-test. ns: non-significant. The number of analyzed mice is indicated in red.

**Supplementary Fig. 10. CC triggers spine morphology alterations in *ex vivo* organotypic hippocampal slice cultures.** (**A**) Representative images of organotypic hippocampal slices (scale bars, 1 μm) and of dendritic segments expressing Green or APPswe-Green mice treated with CC (10 μM) for 6 h. controls are shown in Figure 8. (**B**) Graph representing the percentage of spine types (F: filopodia, I: immature, and M: mature). P values were calculated using Kruskal-Wallis test and Dunn's multiple comparisons post-test (D, F). * $P$ <0.05, ** $P$ <0.01, *** $P$ <0.001, **** $P$ <0.0001 and ns: non-significant. # $P$ <0.05 is relative to spine type in the Green NT condition.

**References**

1. Oules B, Del Prete D, Greco B, Zhang X, Lauritzen I, Sevalle J, et al. Ryanodine receptor blockade reduces amyloid-beta load and memory impairments in Tg2576 mouse model of Alzheimer disease. J Neurosci. 2012;32(34):11820-34.

2. Mu J, Brozinick JT, Jr., Valladares O, Bucan M, Birnbaum MJ. A role for AMP-activated protein kinase in contraction- and hypoxia-regulated glucose transport in skeletal muscle. Mol Cell. 2001;7(5):1085-94.

3. Vaillant-Beuchot L, Mary A, Pardossi-Piquard R, Bourgeois A, Lauritzen I, Eysert F, et al. Accumulation of amyloid precursor protein C-terminal fragments triggers mitochondrial

structure, function, and mitophagy defects in Alzheimer's disease models and human brains. Acta Neuropathol. 2021;141(1):39-65.

A

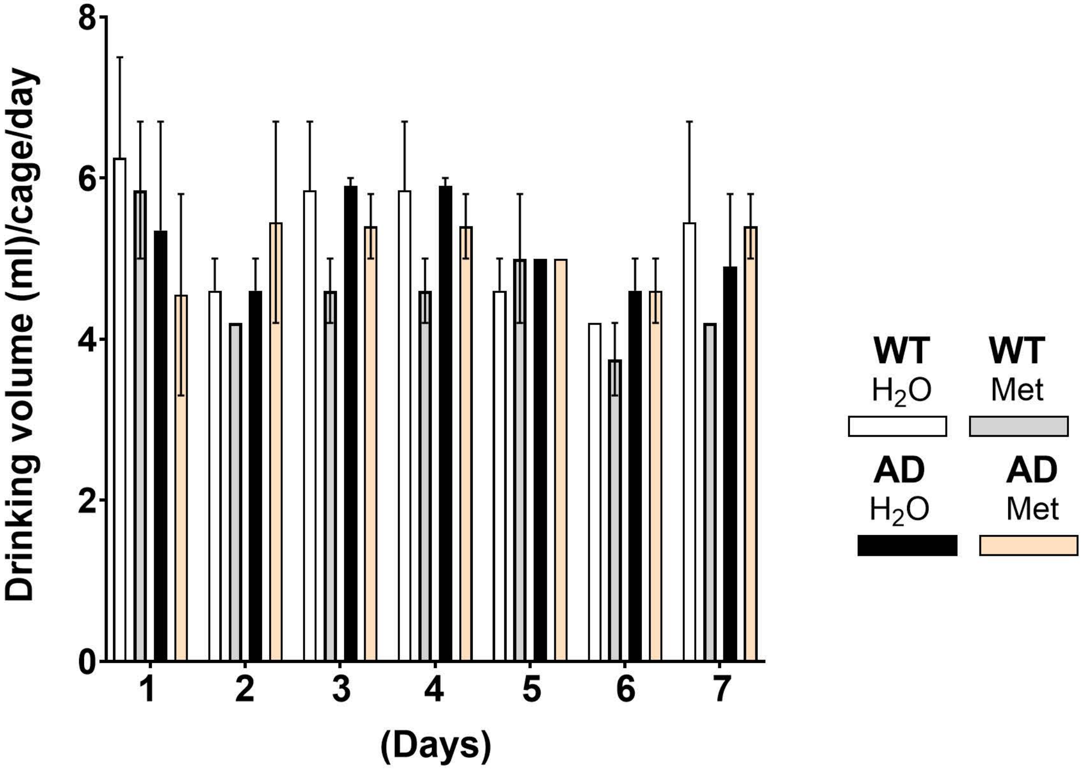

8
6
4
2
0
Drinking volume (ml)/cage/day
1
2
3
4
5
6
7
(Days)
WT
$H_2O$
WT
Met
AD
$H_2O$
AD
Met


B

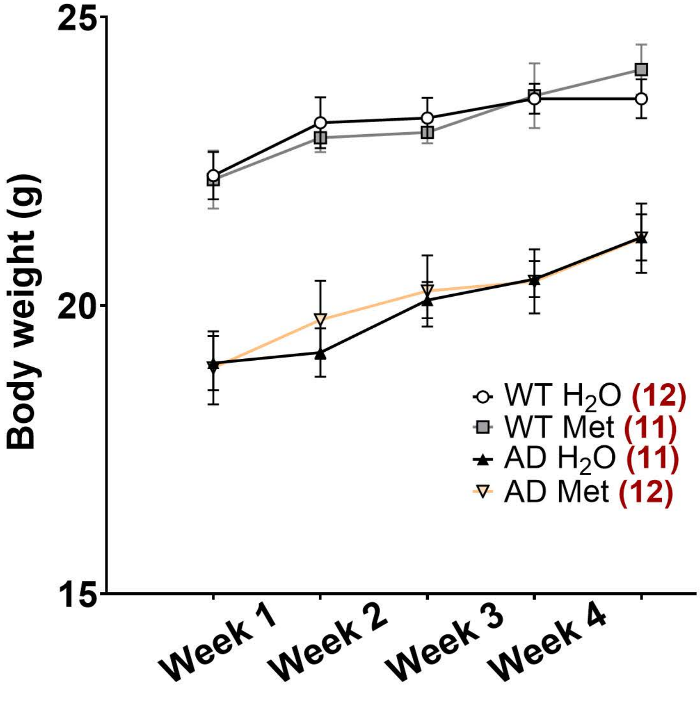

25
20
15
Body weight (g)
Week 1
Week 2
Week 3
Week 4
WT $H_2O$ (12)
WT Met (11)
AD $H_2O$ (11)
AD Met (12)

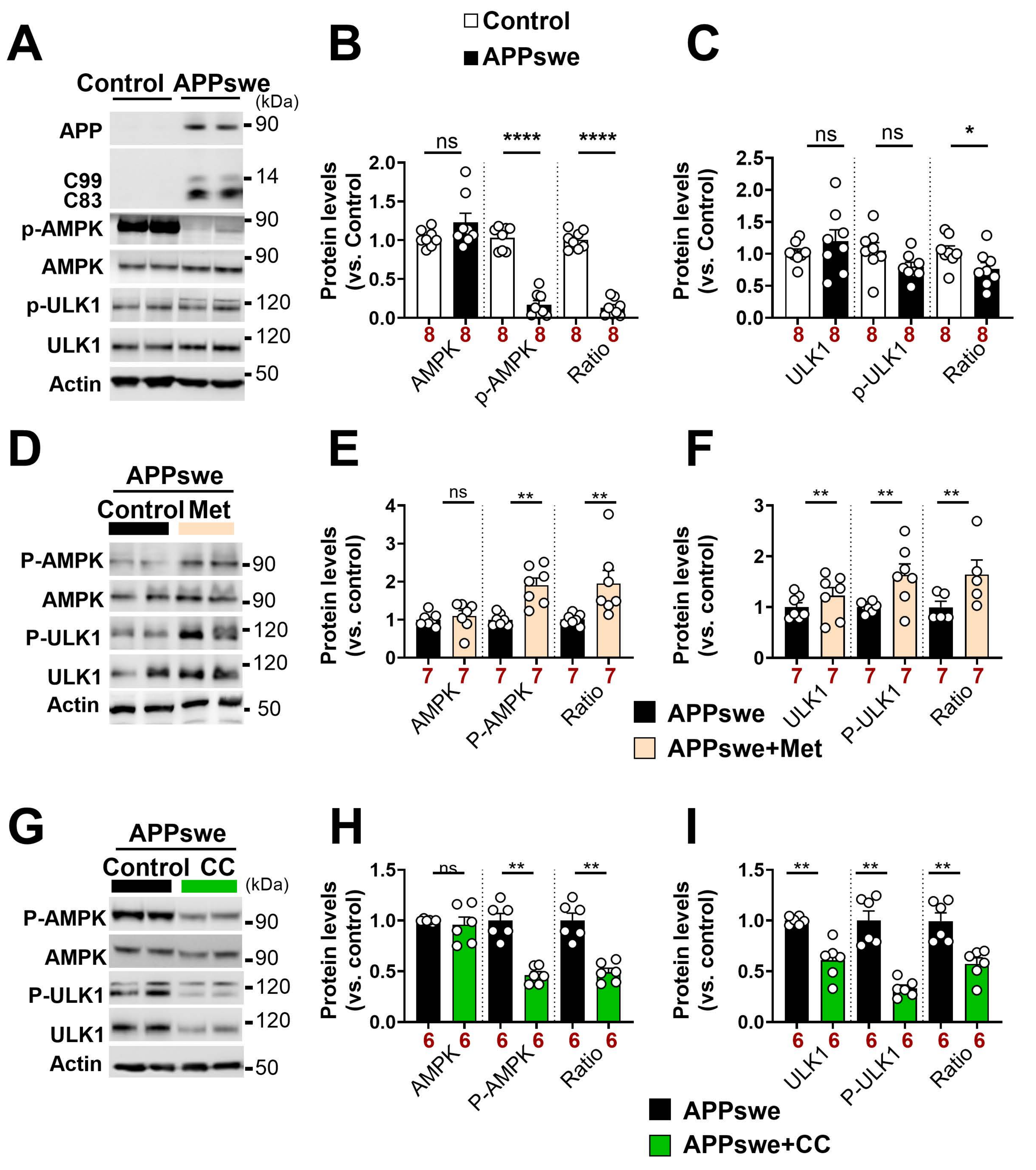

A
Control
APPswe
(kDa)
APP
C99
C83
p-AMPK
AMPK
p-ULK1
ULK1
Actin
90
14
90
90
120
120
50
B
Control
APPswe
Protein levels (vs. Control)
ns
****
****
AMPK
p-AMPK
Ratio
C
Protein levels (vs. Control)
ns
ns
*
ULK1
p-ULK1
Ratio
D
APPswe
Control
Met
P-AMPK
AMPK
P-ULK1
ULK1
Actin
E
Protein levels (vs. control)
ns
**
**
AMPK
P-AMPK
Ratio
F
Protein levels (vs. control)
**
**
**
ULK1
P-ULK1
Ratio
APPswe
APPswe+Met
G
APPswe
Control
CC
(kDa)
P-AMPK
AMPK
P-ULK1
ULK1
Actin
H
Protein levels (vs. control)
ns
**
**
AMPK
P-AMPK
Ratio
I
Protein levels (vs. control)
**
**
**
ULK1
P-ULK1
Ratio
APPswe
APPswe+CC

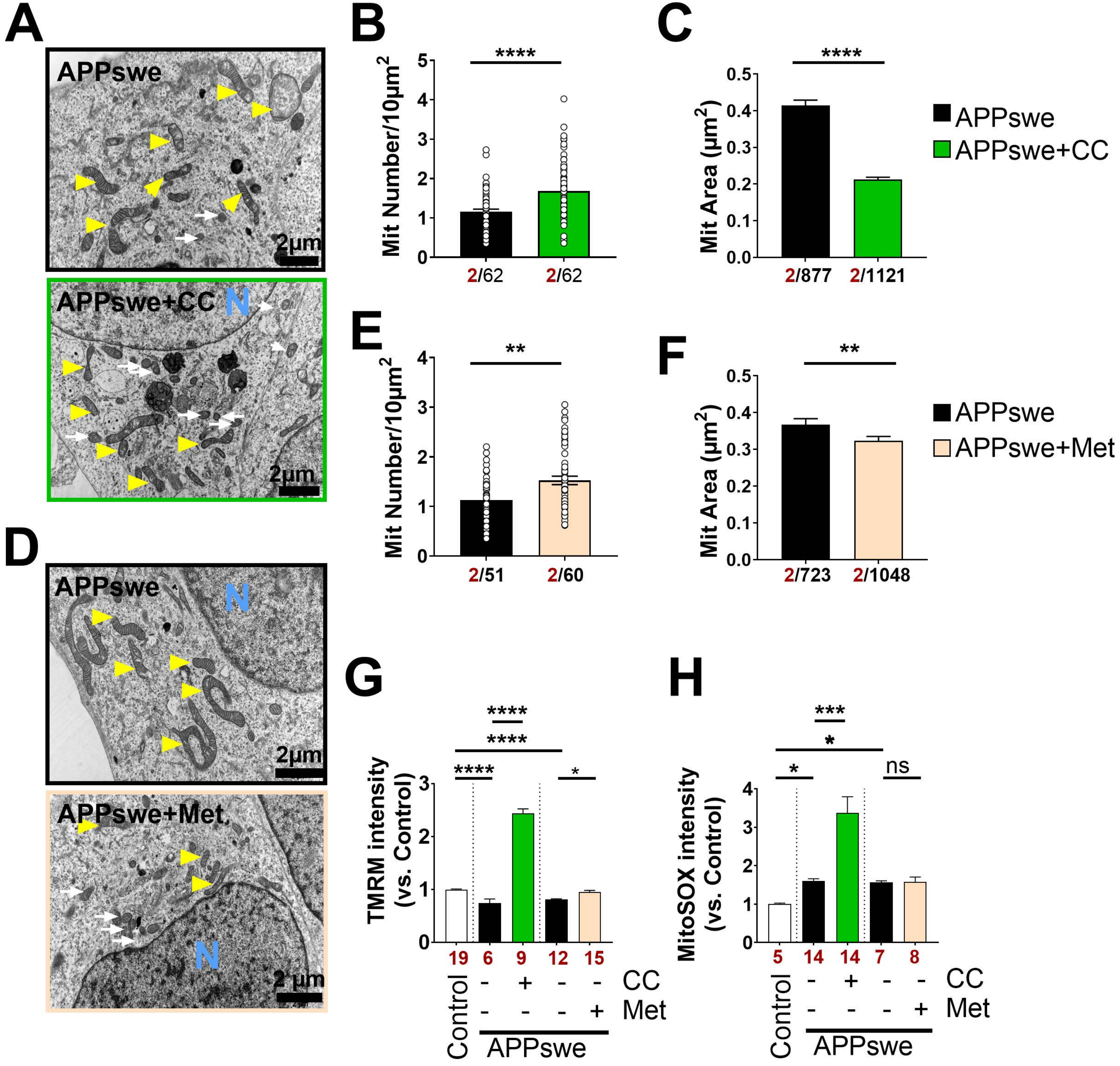

A
APPswe
2μm
APPswe+CC
N
2μm
B
****
Mit Number/10μm2
2/62
2/62
C
****
Mit Area (μm2)
2/877
2/1121
APPswe
APPswe+CC
D
APPswe
N
2μm
APPswe+Met
N
2 μm
E
**
Mit Number/10μm2
2/51
2/60
F
**
Mit Area (μm2)
2/723
2/1048
APPswe
APPswe+Met
G
****
****
****
*
TMRM intensity (vs. Control)
19 6 9 12 15
CC
Met
Control
APPswe
H
***
*
*
ns
MitoSOX intensity (vs. Control)
5 14 14 7 8
CC
Met
Control
APPswe

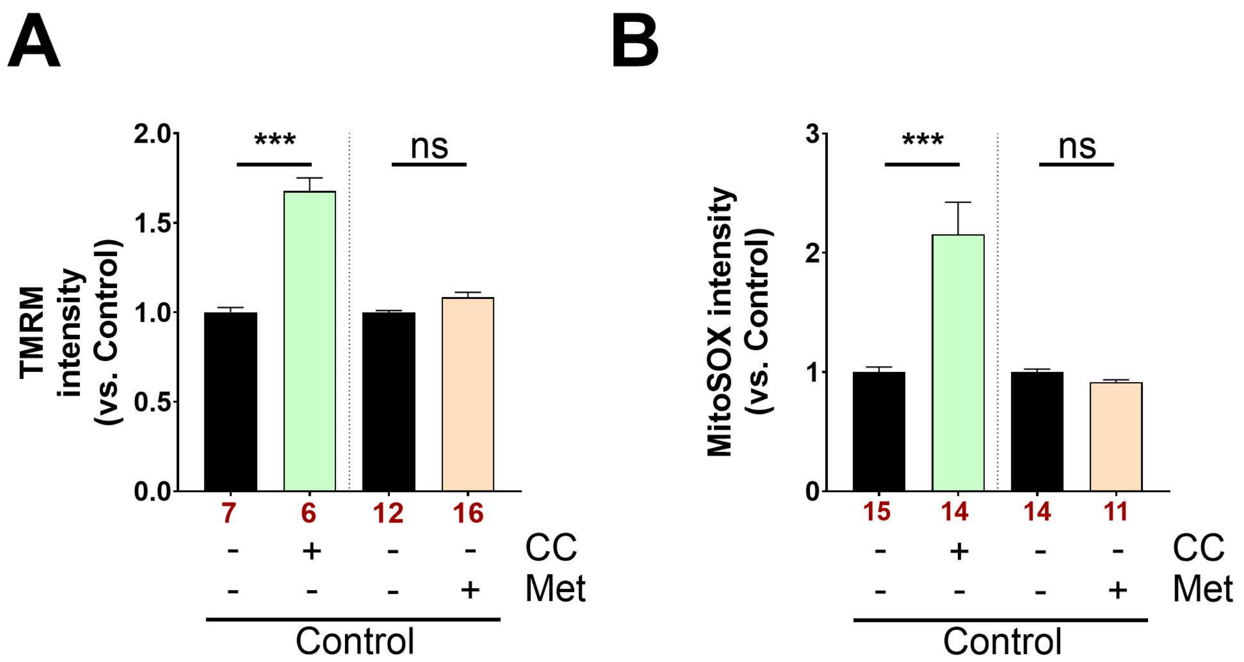
A
B
TMRM intensity (vs. Control)
2.0
1.5
1.0
0.5
0.0
***
ns
7
6
12
16
CC
Met
Control
MitoSOX intensity (vs. Control)
3
2
1
0
15
14
14
11

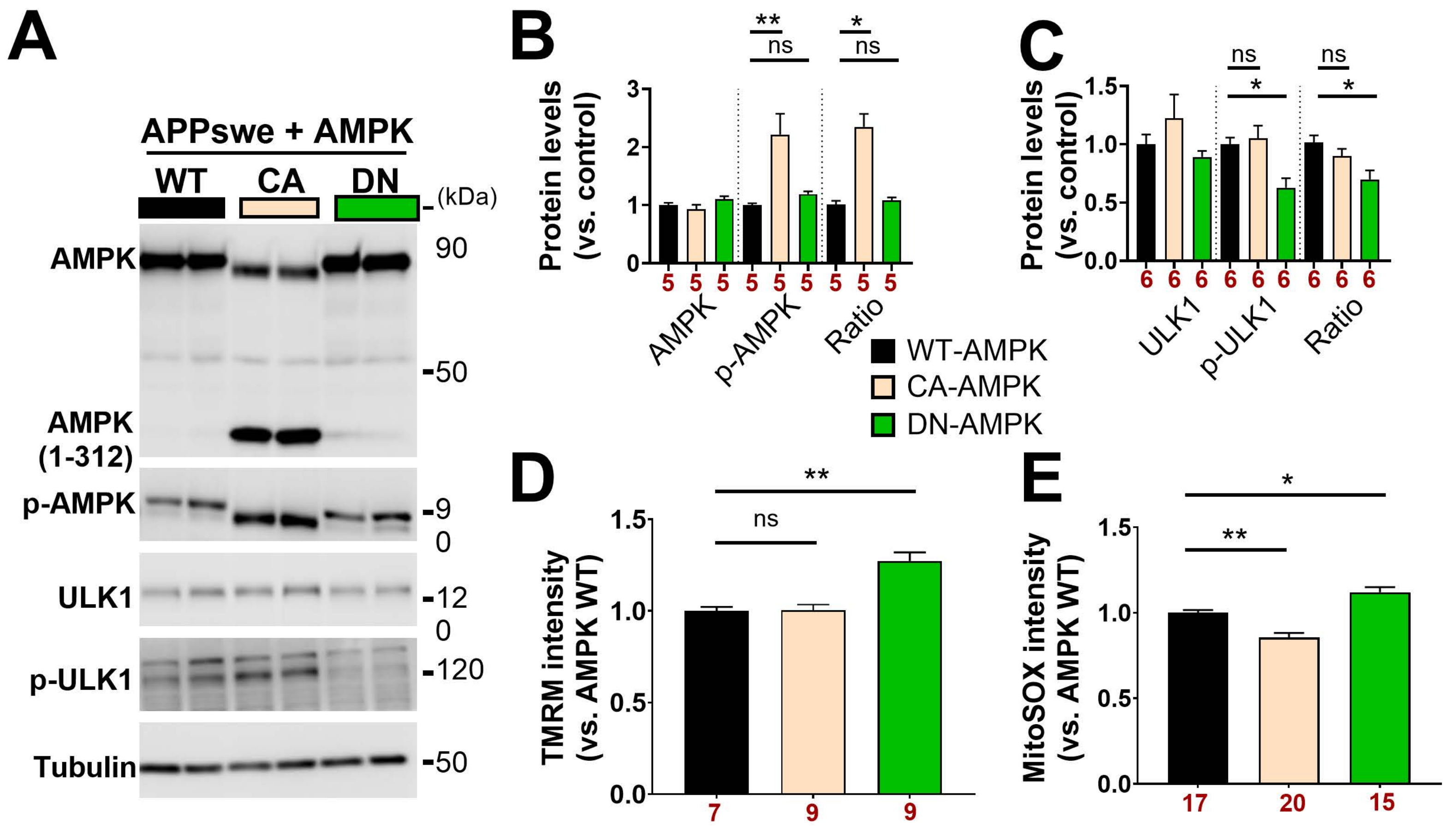
A
APPswe + AMPK
WT
CA
DN
(kDa)
AMPK
90
-50
AMPK (1-312)
p-AMPK
-90
ULK1
-120
p-ULK1
-120
Tubulin
-50
B
Protein levels (vs. control)
AMPK
p-AMPK
Ratio
**
ns
*
ns
5 5 5 5 5 5 5 5 5
C
Protein levels (vs. control)
ULK1
p-ULK1
Ratio
ns
*
ns
*
6 6 6 6 6 6 6 6 6
WT-AMPK
CA-AMPK
DN-AMPK
D
TMRM intensity (vs. AMPK WT)
**
ns
7 9 9
E
MitoSOX intensity (vs. AMPK WT)
*
**
17 20 15

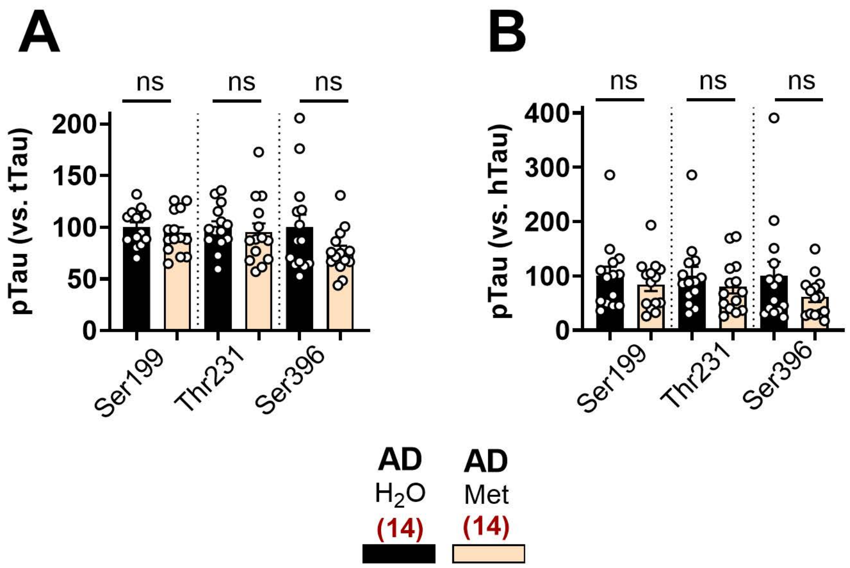

A
B
ns
ns
ns
pTau (vs. tTau)
200
150
100
50
0
Ser199
Thr231
Ser396
pTau (vs. hTau)
400
300
200
100
0
AD
H2O
(14)
AD
Met
(14)

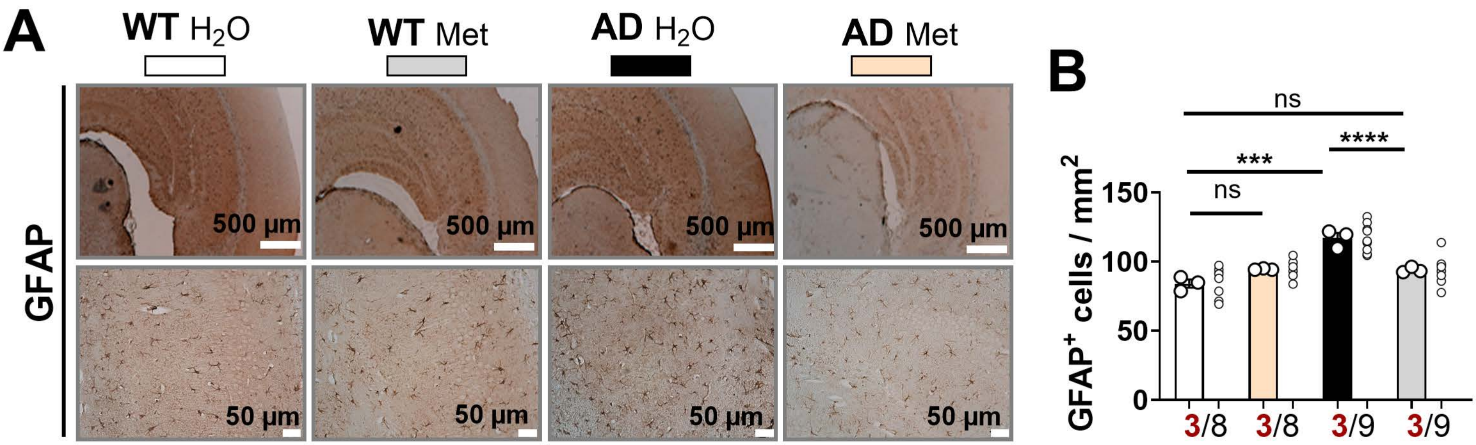
A
WT H2O
WT Met
AD H2O
AD Met
GFAP
500 µm
500 µm
500 µm
500 µm
50 µm
50 µm
50 µm
50 µm
B
GFAP+ cells / mm2
150
100
50
0
ns
***
****
ns
3/8
3/8
3/9
3/9

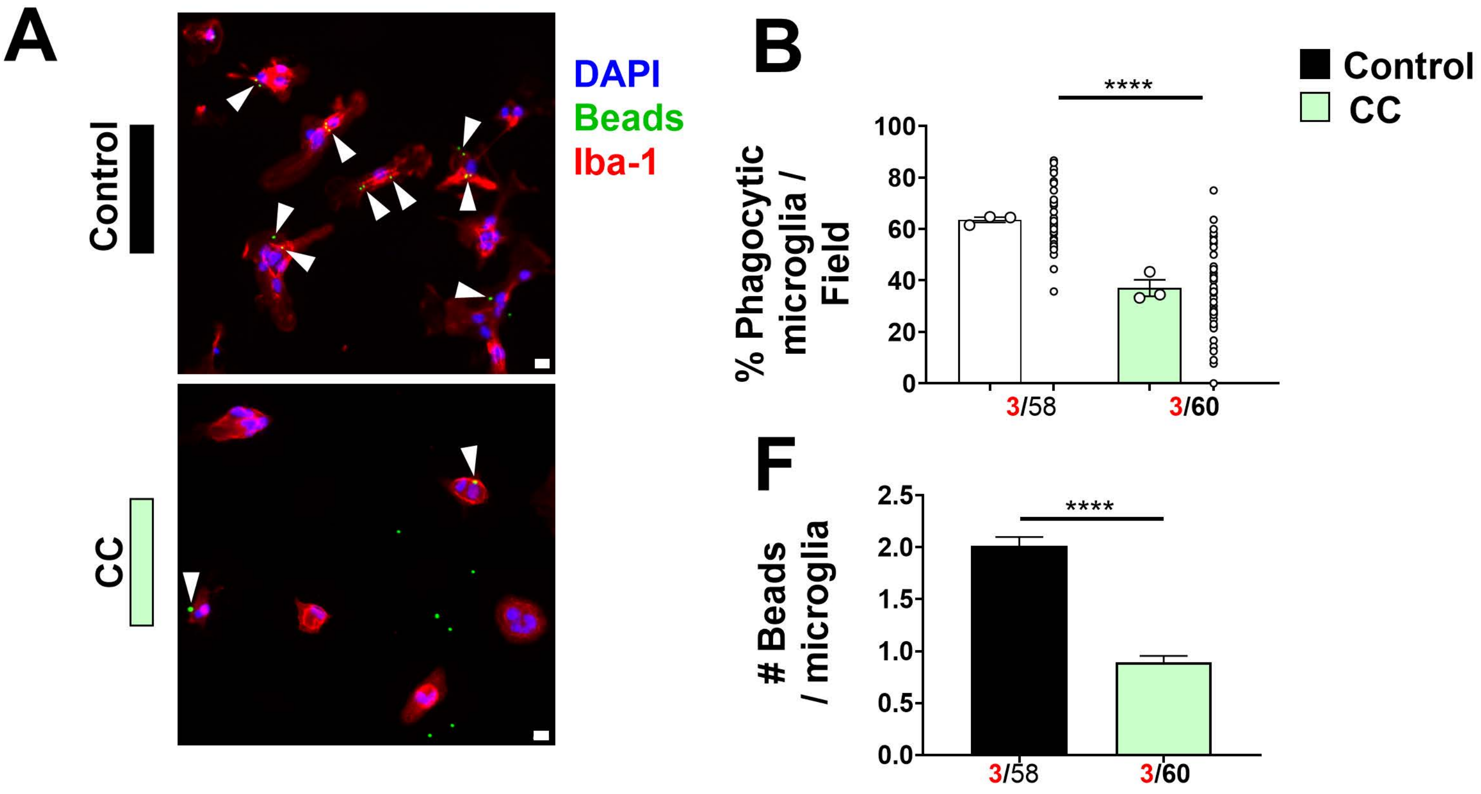
A
Control
CC
DAPI
Beads
Iba-1
B
Control
CC
****
% Phagocytic microglia / Field
100
80
60
40
20
0
3/58
3/60
F
****
### Beads / microglia
2.5
2.0
1.5
1.0
0.5
0.0
3/58
3/60

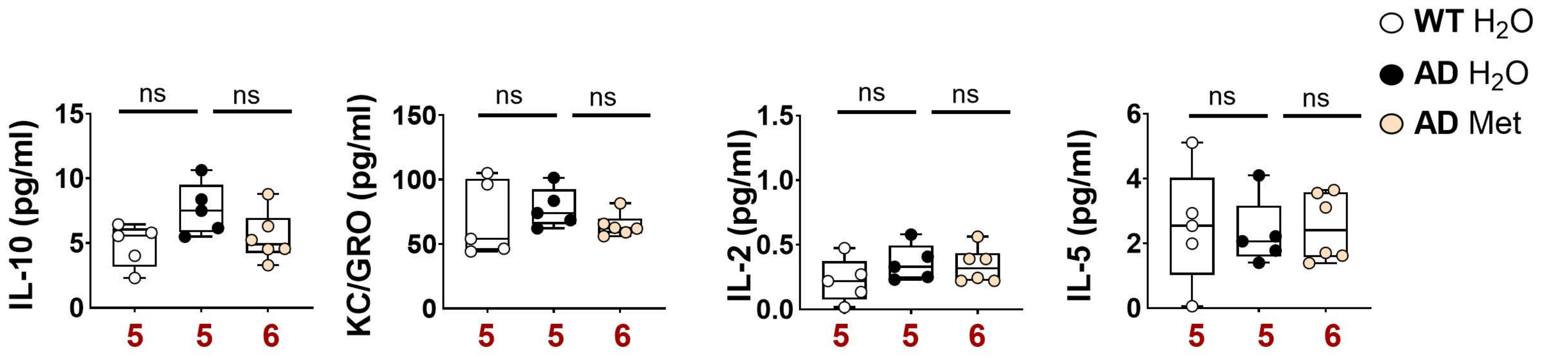

WT H₂O
AD H₂O
AD Met
IL-10 (pg/ml)
KC/GRO (pg/ml)
IL-2 (pg/ml)
IL-5 (pg/ml)
ns
5
6

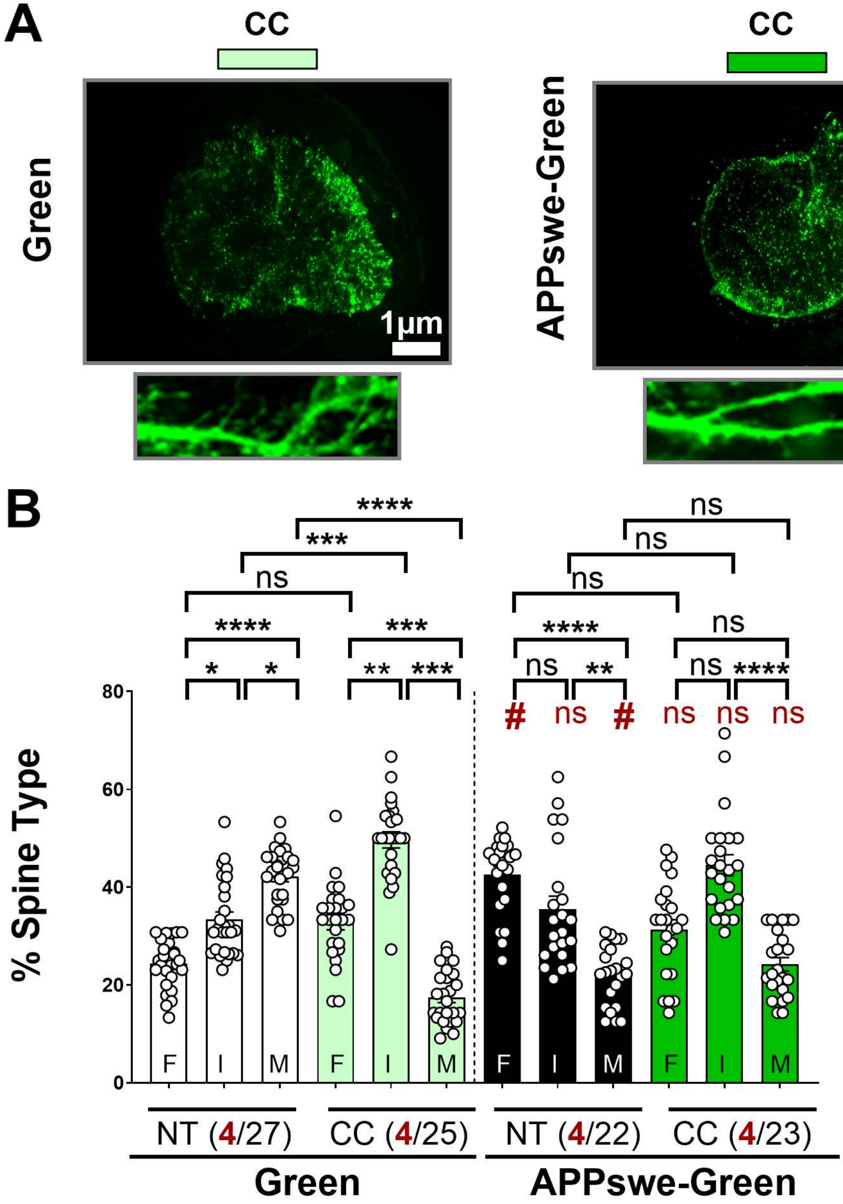
A
CC
CC
Green
APPswe-Green
1µm
1µm
B
% Spine Type
0
20
40
60
80
****
***
ns
****
*
*
***
**
***
ns
ns
ns
****
ns
**
ns
ns
****
#
ns
#
ns
ns
ns
F
I
M
F
I
M
F
I
M
F
I
M
NT (4/27)
CC (4/25)
NT (4/22)
CC (4/23)
Green
APPswe-Green
